# PARK15/FBXO7 is dispensable for PINK1/Parkin mitophagy in iNeurons and HeLa cell systems

**DOI:** 10.1101/2022.11.02.514817

**Authors:** Felix Kraus, Ellen A. Goodall, Ian R. Smith, Yizhi Jiang, Julia C. Paoli, Frank Adolf, Jiuchun Zhang, Joao A. Paulo, Brenda A. Schulman, J. Wade Harper

## Abstract

The protein kinase PINK1 and ubiquitin ligase Parkin promote removal of damaged mitochondria via a feed-forward mechanism involving ubiquitin (Ub) phosphorylation (pUb), Parkin activation, and ubiquitylation of mitochondrial outer membrane proteins to support recruitment of mitophagy receptors. The ubiquitin ligase substrate receptor FBXO7/PARK15 is mutated in an early-onset parkinsonian-pyramidal syndrome. Previous studies have proposed a role for FBXO7 in promoting Parkin-dependent mitophagy. Here, we systematically examine the involvement of FBXO7 in depolarization and ^mt^UPR-dependent mitophagy in the well-established HeLa and induced-neurons cell systems. We find that FBXO7^-/-^ cells have no demonstrable defect in: 1) kinetics of pUb accumulation, 2) pUb puncta on mitochondria by super-resolution imaging, 3) recruitment of Parkin and autophagy machinery to damaged mitochondria, 4) mitophagic flux, and 5) mitochondrial clearance as quantified by global proteomics. Moreover, global proteomics of neurogenesis in the absence of FBXO7 reveals no obvious alterations in mitochondria or other organelles. These results argue against a general role for FBXO7 in Parkin-dependent mitophagy and point to the need for additional studies to define how FBXO7 mutations promote parkinsonian-pyramidal syndrome.

## Introduction

Organelle quality control underlies cellular health and is often defective in disease and pathological states. Arguably the best understood such quality control pathway is mitophagy, wherein damaged or supernumerary mitochondria are targeted for removal from the cell via selective autophagy (Goodall *et al*, 2022; Harper *et al*, 2018; McWilliams & Muqit, 2017; Pickrell & Youle, 2015). In this process, individual organelles are marked for sequestration within a double-membrane vesicle called an autophagosome, which then fuses with a lysosome to facilitate degradation of the organelle by resident lysosomal hydrolases (Dikic & Elazar, 2018; Stolz *et al*, 2014). Multiple forms of mitophagy have been reported, which differ in the types of regulatory mechanisms involved in marking the organelle for degradation, but these fall into two primary types – ubiquitin-dependent and ubiquitin-independent (Harper *et al*., 2018; Pickrell & Youle, 2015). Our understanding of ubiquitin-dependent mitophagy has been advanced by the discovery of a signalling pathway composed of the PINK1 protein kinase and the Parkin (also called PRKN) ubiquitin (Ub) E3 ligase, which marks damaged mitochondria for elimination (Narendra *et al*, 2008). Parkin and PINK1 are both mutated in early-onset recessive forms of Parkinson’s disease and understanding how these enzymes work has been a major focus of the field (Ng *et al*, 2021; Pickrell & Youle, 2015).

In healthy mitochondria, PINK1 is imported into the mitochondrial translocon and rapidly processed for degradation (Jin *et al*, 2010; Yamano & Youle, 2013). In response to mitochondrial damage – such as depolarization, accumulation of mitochondrial misfolded proteins, or defects in mitochondrial fusion – PINK1 is stabilized on the mitochondrial outer membrane (MOM) in association with the translocon (Lazarou *et al*, 2012), where it can phosphorylate Ser65 on Ub already conjugated to proteins on the MOM (Kane *et al*, 2014; Kazlauskaite *et al*, 2015; Koyano *et al*, 2014; Ordureau *et al*, 2014; Wauer *et al*, 2015). Accumulation of phospho S65-Ub (referred to henceforth as pUb) on the MOM promotes recruitment of Parkin – a pUb-binding protein – thereby facilitating phosphorylation of Parkin on S65 of its N-terminal Ub-like (UBL) domain and activation of its Ub ligase activity (Gladkova *et al*, 2018; Kane *et al*., 2014; Kazlauskaite *et al*., 2015; Ordureau *et al*., 2014; Sauve *et al*, 2018; Wauer *et al*., 2015). Parkin then ubiquitylates numerous MOM proteins, resulting in both the accumulation of additional pUb and further MOM protein ubiquitylation (Antico *et al*, 2021; Bingol *et al*, 2014; Ordureau *et al*, 2020; Sarraf *et al*, 2013). MOM ubiquitylation then promotes recruitment of selective autophagy cargo receptors including OPTN, CALCOCO2 (also called NDP52), and SQSTM1 (also called p62) to facilitate the assembly of an autophagosome around the ubiquitylated organelle (Evans & Holzbaur, 2020; Heo *et al*, 2015; Lazarou *et al*, 2015; Wong & Holzbaur, 2014). Thus, PINK1, Parkin, and pUb function in a positive feedback loop to promote selective ubiquitylation and elimination of the damaged organelle (Goodall *et al*., 2022). Removal of damaged mitochondria may be an important aspect of neuronal protection in both neurodegenerative diseases and in aging, and understanding how the PINK1/Parkin pathway functions in this regard is a central goal of the field (Fang *et al*, 2019; Fleming *et al*, 2022).

Previous work (Burchell *et al*, 2013) concluded that FBXO7, the product of an early-onset parkinsonian-pyramidal syndrome risk gene also designated as *PARK15* (Di Fonzo *et al*, 2009; Houlden & Singleton, 2012; Paisan-Ruiz *et al*, 2010), is a positive modulator of Parkin-dependent mitophagy (**Appendix Figure S1A**). FBXO7 is a member of the F-box family of proteins, which assemble with SKP1, CUL1, and RBX1 to form a modular SCF Ub ligase complex wherein the F-box protein binds substrates (Jin *et al*, 2004). FBXO7 is characterized by an N-terminal UBL domain, a central FP domain with structural similarity to a domain in the proteasome inhibitory factor PI31/PSMF1, followed by the F-box motif, and a C-terminal proline rich region (Kirk *et al*, 2008). Patient mutations in FBXO7 are found in multiple regions of the protein (Di Fonzo *et al*., 2009; Houlden & Singleton, 2012; Paisan-Ruiz *et al*., 2010). Initial studies on FBXO7 proposed a functional interaction with the PINK1/Parkin pathway and concluded that FBXO7 functions as a positive regulator of mitophagy through direct interaction with Parkin and with PINK1 (Burchell *et al*., 2013). Depletion of FBXO7 by siRNA in cell lines was reported to reduce depolarization-dependent loss in mitochondria based on immunoblotting of matrix proteins and likewise to reduce depolarization-dependent ubiquitylation of MFN1, a substrate of Parkin (Burchell *et al*., 2013). However, studies that further substantiate a functional link between FBXO7 and mitophagy are lacking, although FBXO7 depletion has recently been reported to alter mitochondrial dynamics or PINK1 levels (Al Rawi *et al*, 2022; Huang *et al*, 2020; Liu *et al*, 2020).

Given the strong genetic association of FBXO7/PARK15 with Parkinson’s Disease (Houlden & Singleton, 2012) and major advances in our understanding of Parkin-dependent mitophagy (Goodall *et al*., 2022) since initial links between FBXO7 and mitophagy were reported, we set out to elucidate how FBXO7 might function within the positive feedback loop to promote mitophagy (**Appendix Figure S1A**). Relevant advances include the identification of pUb as a sentinel response to PINK1/Parkin activation (Kane *et al*., 2014; Kazlauskaite *et al*., 2015; Koyano *et al*., 2014; Ordureau *et al*., 2014; Wauer *et al*., 2015), the development of highly quantitative mitophagic flux assays using a mitochondrially localized Keima fluorescent reporter protein (Katayama *et al*, 2011; Lazarou *et al*., 2015), and the development of genetically tractable iNeuron systems for functional analysis of the Parkin pathway in neuronal cell types (Ordureau *et al*., 2020; Ordureau *et al*, 2018). Unexpectedly, we find no evidence of a role for FBXO7 in the kinetics of depolarization-dependent pUb accumulation, or mitophagic flux in either iNeurons or the conventional HeLa cell used extensively to examine PINK1/Parkin-dependent mitophagy (Narendra *et al*., 2008). Moreover, global proteomics of FBXO7^-/-^ HeLa cells undergoing mitophagy revealed no defect in mitochondrial elimination by autophagy. Taken together with previous studies that failed to verify Parkin-FBXO7 interactions during mitophagy (Sarraf *et al*., 2013), these data indicate that FBXO7 does not play a general role as a positive modulator of PINK1/Parkin-dependent mitophagy in HeLa and iNeuron systems. This work suggests that elucidation of FBXO7 biochemical functions is needed to understand how it might contribute to suppression of early-onset parkinsonian-pyramidal syndrome.

## Results

### A toolkit for functional analysis of cells lacking FBXO7

To test if FBXO7 is a pivotal amplifier of the PINK1/Parkin pathway, we employed CRISPR-Cas9-based gene editing to generate knockouts for FBXO7 in HeLa HFT/TO-PRKN cells (Ordureau *et al*, 2015) as well as HEK293T cells and confirmed insertion of a frameshift mutation by sequencing and immunoblotting (**Appendix Figure S1C-F**). HeLa HFT/TO-PRKN cells contain a DOX-inducible PRKN cassette allowing regulated expression of Parkin to facilitate an analysis of mitophagic flux in response to mitochondrial damage (Ordureau *et al*., 2015). Three independent HeLa HFT/TO-PRKN FBXO7^-/-^ clones were identified (clones 27, 41 and 45) (**Appendix Figure S1C, D**). Two independent HEK293T cell lines lacking FBXO7 were transduced with a lentivirus for stable expression of GFP-Parkin to facilitate pathway activation. In parallel, we generated two independent human ES cell clones lacking FBXO7 via CRISPR-Cas9 (clones 52 and 89) (**Appendix Figure S1G-I**). Here, an ES cell line harbouring a DOX-inducible NGN2 gene was used, allowing for conversion of these cells to cortical-like iNeurons over a two week or greater time course (Ordureau *et al*., 2020; Zhang *et al*, 2013). We have previously demonstrated that iNeurons derived from these cells exhibit PINK1/Parkin-dependent mitophagic flux in response to mitochondrial damage (Ordureau *et al*., 2020). Both clones displayed normal karyotypes and the absence of FBXO7 was verified using MiSeq sequencing (**Appendix Figure S1 H, I**).

### Robust depolarization-dependent Ub phosphorylation by PINK1 in cells lacking FBXO7

The earliest known event in Parkin-dependent mitophagy is accumulation of PINK1 on the mitochondrial outer membrane followed by rapid S65 phosphorylation of pre-existing Ub linked with mitochondrial proteins (**Appendix Figure S1A**), as has been worked out in detail using the well-validated HeLa cell system with ectopically expressed Parkin (Kane *et al*., 2014; Lazarou *et al*., 2012; Narendra *et al*, 2010; Yamano & Youle, 2013). In the presence of Parkin, Ub phosphorylation is enhanced as a result of increased Ub conjugation to mitochondrial proteins, thereby providing additional substrate molecules for phosphorylation by PINK1 (Kane *et al*., 2014; Koyano *et al*., 2014; Ordureau *et al*., 2014). To examine pUb accumulation in the context of HeLa cells lacking FBXO7, we induced Parkin expression (16 h) and then depolarized mitochondria for 1 or 6 h with antimycin A/Oligomycin A (AO), which inhibit Complex III and Complex IV activity in the electron transport chain, respectively. As expected, control HeLa cells display time-dependent accumulation of pUb as measured by immunoblotting, but the absence of FBXO7 had no impact on the extent of pUb accumulation, as assessed in all three FBXO7^-/-^ cell lines in biological triplicate analyses (**Figure 1A,B**). Additionally, we used CCCP (Carbonyl cyanide m-chlorophenyl hydrazone) as an alternative to induce mitophagy and found pUb levels in either HeLa and HEK293T cells were similar in control and FBXO7^-/-^ cells (**Appendix Figure S1J,K**).

**Figure 1:**
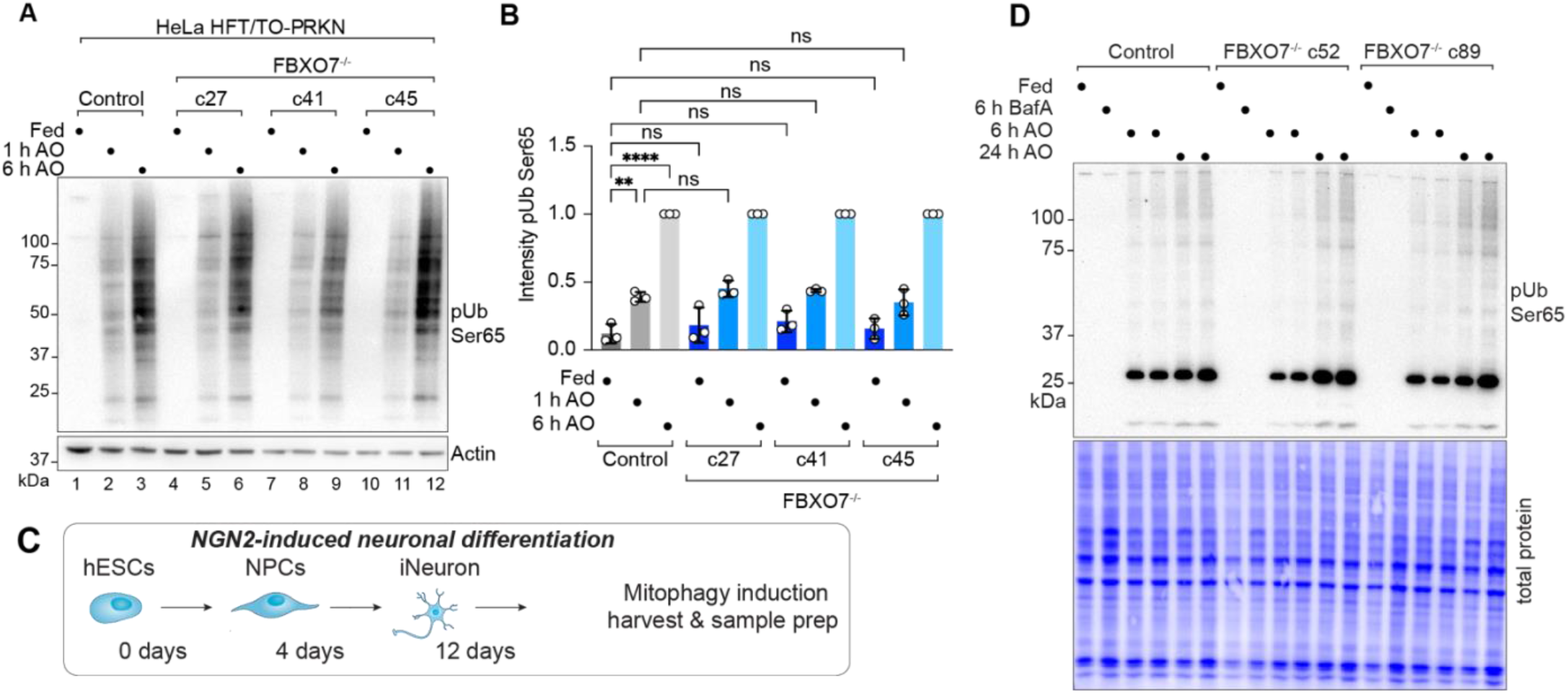
Robust depolarization-dependent Ub phosphorylation by PINK1 cells lacking FBXO7. (**A**) Immunoblot for pUb on HeLa control and FBXO7^-/-^ cells after treatment with AO for indicated times. (**B**) Quantification of pUb signal relative to loading. Immunoblots run in triplicates. Error bars depict S.D.. Data from three biological replicates. One-way ANOVA with multiple comparisons. p(**) = 0.0011; p(****)<0.0001. (**C**) Scheme depicting production of iNeurons for biochemical analysis. (**D**) Immunoblot for pUb on extracts from day 12 iNeuron control and FBXO7 -^/-^ cells after treatment with BafA or AO for indicated times.

Similarly, we assessed depolarization-dependent pUb accumulation in the iNeuron system. control or FBXO7^-/-^ ES cells were converted to day 12 iNeurons and depolarized with AO in duplicate (**Figure 1C**). pUb was detected by immunoblotting at 6 and 24 h and was indistinguishable between WT and two independent FBXO7^-/-^ clones (**Figure 1D**). Taken together, these data do not support a universal role for FBXO7 in promoting the earliest step in mitophagy signalling – PINK1-dependent Ub phosphorylation – in the HeLa cell system or iNeurons.

### Super-resolution imaging of pUb formation independent of FBXO7

While depolarization-dependent Ub phosphorylation was not impaired, we considered the possibility that the organization of pUb on damaged mitochondria might be altered in cells lacking FBXO7. To our knowledge, super-resolution microscopy has not been used to examine the kinetics and spatial properties of pUb accumulation in response to depolarization. We therefore optimized conditions to monitor pUb via volumetric 3D-SIM super-resolution microscopy in both HeLa and iNeuron systems (see **Materials and Methods**). After AO treatment of HeLa cells for 1, 3 or 6 h, the mitochondria matrix protein HSP60 and pUb were examined by super-resolution imaging. In control cells at 1 h, fragmented mitochondria were often proximal to pUb foci with an average volume of 5.71 μm^3^, which were not observed in cells lacking PINK1 (**Figure 2A-C**, **Figure EV 1A,B**). The volume of pUb increased ∼8-fold over 6 h (47.88 μm^3^), consistent with the known feed-forward response (**Figure 2B, Figure EV 1A**). At 6 h of AO treatment, we observed a near complete coating of mitochondrial fragments with pUb, as quantified by the significant increase in pUb object volume (**Figure 2B,C, Figure EV 1A**). Consistent with analysis of pUb by immunoblotting, two FBXO7^-/-^ clones did not show any significant differences in pUb recruitment, volume or overall morphological differences compared to control cells, indicating that FBXO7 is not required for the feed-forward amplification of PINK1/Parkin mitophagy after AO-treatment (**Figure 2A-C**). Analogous super-resolution experiments in day 12 iNeurons with 0.5, 1, and 3 h of AO revealed similar results, with no obvious effect of FBXO7 deletion on pUb recruitment, volume or overall morphology when compared to WT iNeurons (**Figure 2D-F, Appendix Figure S2A**). As expected (Ordureau *et al*., 2020; Ordureau *et al*., 2018), iNeurons lacking PINK1 did not accumulate pUb (**Figure 2D,E**). These results were also confirmed in iNeurons treated with AO for 1 or 6 h using conventional immunofluorescence and image analysis of HSP60 and pUb co-localization (**Figure 2G,H**). To ensure that kinetic effects were not obscured by overt depolarization, we reduced the concentrations of AO used by ∼10-fold, but again no defect in pUb recruitment to mitochondria was observed (**Appendix Figure S2B,C**). These data indicate that the absence of FBXO7 does not alter the ability of PINK1-dependent pUb foci to accumulate on damaged mitochondria in either HeLa or iNeuron system

**Figure 2.**
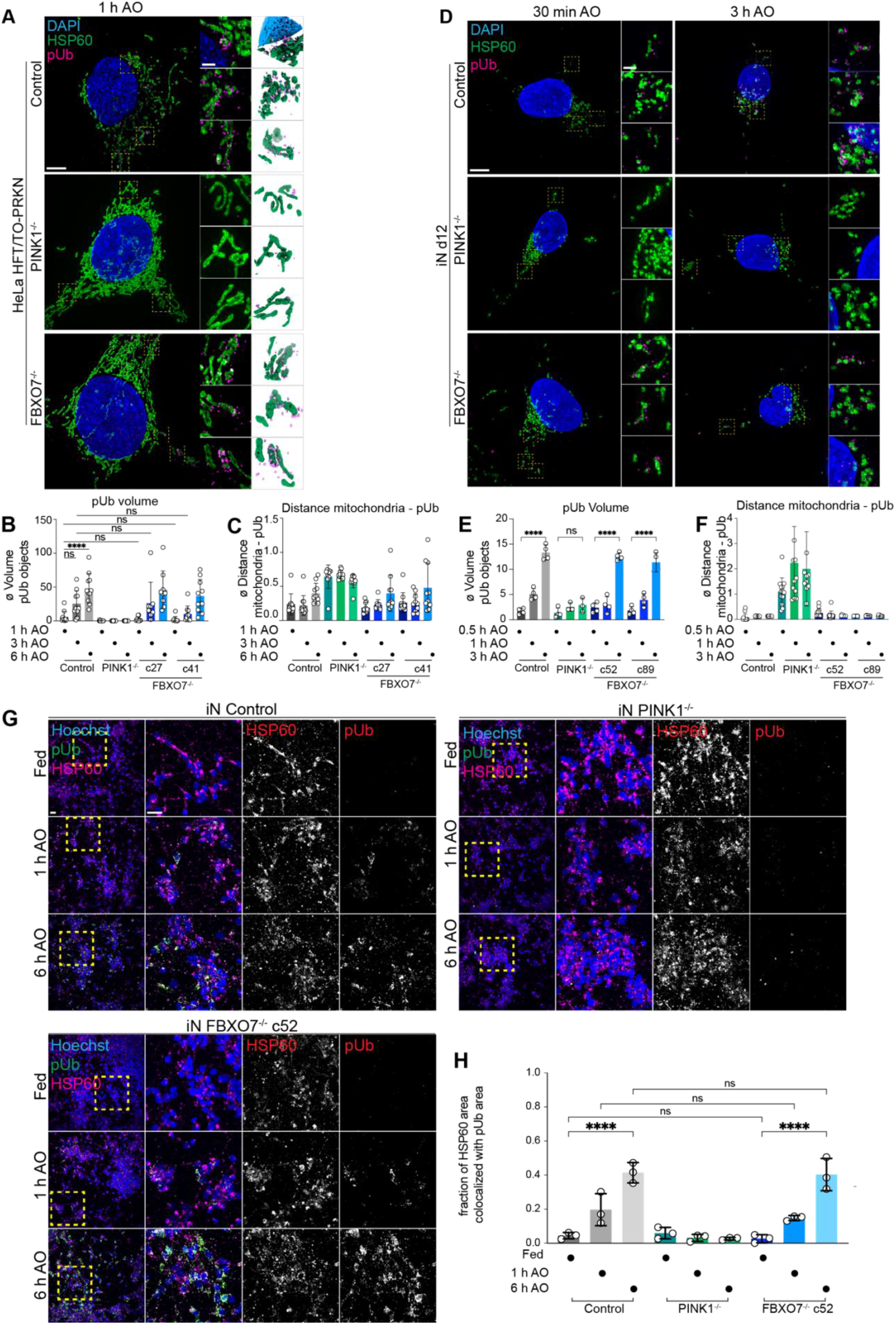
Super-resolution imaging of pUb formation independent of FBXO7. (**A**) 3D-SIM images of HeLa control, PINK1^-/-^ and FBXO7^-/-^ cell lines after AO-induced mitophagy. Cells were stained for nuclear DNA (DAPI), mitochondria (HSP60) and pUb. Zoom-ins of regions of interested are enlarged in the middle panel. 3D-surface renderings of insets are shown on the right. Scale bar = 5 μm or 1 μm. (**B,C**) Evaluation of 3D-SIM images from HeLa datasets. The changes in pUb volume and minimal distances between mitochondria and pUb after mitophagy-induction are plotted. Error bars depict S.D. from 8-14 measured cells per condition. Two-way ANOVA with multiple comparisons; p(****)<0.0001. (**D**) 3D-SIM images of iN day 12 control, PINK1^-/-^ and FBXO7^-/-^ cell lines after AO-induced mitophagy. Cells were stained for nuclear DNA (DAPI), mitochondria (HSP60) and pUb. Zoom-ins of regions of interest are enlarged in the middle panel. 3D-surface renderings of insets are shown on the right. Scale bar = 5 μm or 1 μm. (**E,F**) Evaluation of 3D-SIM images from iNeuron datasets. The changes in pUb volume and minimal distances between mitochondria and pUb after mitophagy-induction are plotted. Error bars depict S.D. from 7-14 measured cells per condition. One-way ANOVA with multiple comparisons, p(****)<0.0001. (**G**) Confocal images of iNeuron d12 Control, PINK1^-/-^ and FBXO7^-/-^ cell lines after AO-induced mitophagy. Cells were stained for nuclear DNA (Hoechst33342), mitochondria (HSP60) and pUb. Scale bar = 10μm and 5μm. (**H**) Evaluation of pUb volume after mitophagy induction. Error bars depict S.D. from 3 replicates. One-way ANOVA with multiple comparisons; p(****)<0.0001.

### Parkin and autophagy regulator recruitment in FBXO7^-/-^ cells

Parkin recruitment depends upon the accumulation of pUb on damaged mitochondria (**Appendix Figure S1A**) (Okatsu *et al*, 2015). Given that the existing model for FBXO7 in Parkin-dependent mitophagy posited that Parkin and FBXO7 physically interact and that FBXO7 promotes Parkin function (Burchell *et al*., 2013), we directly tested whether FBXO7 is required for Parkin recruitment to mitochondria using the HeLa cell system. Control or FBXO7^-/-^ cells were stably transduced with a lentivirus expressing GFP-Parkin and subjected to imaging with or without depolarization with AO (**Figure EV 2A**). We used CellProfiler (Stirling *et al*, 2021) to analyse the characteristics of GFP-Parkin and mitochondria and their relationship to each other. By segmenting out cytosolic and mitochondrially-localized Parkin, we could quantify alterations in Parkin abundance within the mitochondrial mask as a measure of Parkin-translocation (**Figure EV 2B**). Our expectation was that a requirement for FBXO7 in Parkin recruitment would result in reduced Parkin intensity within the mitochondrial mask in response to depolarization. First, we tested whether mitochondrial morphology (mitochondrial integrated density [==mtIntDen] as proxy, stained with HSP60) is changing after AO-treatment on a single-cell level. Under fed conditions, control and three FBXO7^-/-^ clones displayed comparable mtIntDen. Depolarization by AO led to the expected mitochondrial fragmentation and clustering, thus causing an increase in local HSP60 signal and a right-shift in mtIntDen (**Figure EV 2C, top panel**). This shift was also observed to a similar extent in FBXO7^-/-^ cells. Thus, both control and FBXO7^-/-^ cells robustly respond to AO-induced mitophagy. Next, we tested if Parkin intensities are increasing on mitochondria on an “organelle object” level. As expected, mtIntDen did shift to the right after AO-treatment, in line with previous experiments. Parkin signal was comparable between the genotypes under fed conditions, and increased in intensity after AO-treatment, consistent with Parkin accumulation inside the mitochondrial area (**Figure EV 2C, lower panel**). This Parkin shift was observed in both control and FBXO7^-/-^ cell lines, arguing that FBXO7 is not universally required for Parkin recruitment to damaged mitochondria.

Mitochondrial ubiquitylation by Parkin promotes recruitment of downstream autophagy machinery, including the FIP200-ULK1 complex and Ub-binding autophagy adaptors, ultimately leading to the formation of an autophagosome (Heo *et al*., 2015; Lazarou *et al*., 2015; Ravenhill *et al*, 2019; Vargas *et al*, 2019). Lipidation of LC3 is associated with autophagosome formation and is thought to contribute to recruitment of autophagy receptors (Chang *et al*, 2021). Depolarization of Parkin-expressing HeLa cells resulted in recruitment of both p62 and FIP200 to mitochondria in a PINK1-depenent manner, as revealed by immunostaining and confocal imaging (**Figure EV 2D**). However, FBXO7^-/-^ cells were as proficient as control cells in recruitment of p62 or FIP200 after 3 h or 16 h or AO-induced mitophagy (**Figure EV 2D-G**). Previous studies using siRNAs targeting FBXO7 also reported that LC3 lipidation in response to mitochondrial depolarization is defective compared with non-depleted cells (Burchell *et al*., 2013). However, all three FBXO7^-/-^ HeLa cell clones underwent depolarization-dependent LC3 lipidation to similar extents, consistent with a functional Parkin pathway (**Figure EV 2H,I**). Taken together, these data indicate that FBXO7 is not universally required to promote either Parkin-dependent recruitment to damaged mitochondria or subsequent steps that depend upon Ub chain assembly on mitochondria such as recruitment of autophagy machinery and initiation of LC3 lipidation.

To probe for early mitophagy initiation sites, we treated HeLa control and FBXO7^-/-^ cells with 3h AO or CCCP and stain for the mitochondrial protein Tom20 and WIPI2 (WD repeat domain phosphoinositide-interacting protein 2) (Lazarou *et al*., 2015). Loss of FBXO7 did not lead to a loss of mitophagy initiation, as WIPI2 foci were detectable in Control and FBXO7^-/-^ cell lines (**Figure EV 2J, red arrows**). Evaluation of WIPI2 foci normalized per cell found no statistically difference in foci number between Control and two FBXO7^-/-^ clones after 3h CCCP treatment and a significant difference in only one of the two clones when treated with AO (**Figure EV 2J,K**). Next, we repeated our GFP-Parkin and pUb translocation measurement after mitophagy induction in HEK293T Control and FBXO7^-/-^ cells stably expressing GFP-Parkin. Both independent knockout clones displayed a robust response to 1h AO or CCCP, with both Parkin and pUb accumulating on mitochondria (**Figure EV 2L**). Quantifying the ø pUb intensity showed a significant increase in signal in Control and both knockout clones, supporting the notion that FBXO7 is not essential for PINK1/Parkin mitophagy (**Figure EV 2M**). Taken together, our data from HeLa and HEK293T cell lines suggest that FBXO7 is not essential for the initiation and propagation of mitophagy.

### Interaction network analysis of FBXO7^-/-^

Additionally, we have not been able to reproduce a previously reported association between overexpressed FBXO7 and PINK1 (Huang *et al*., 2020) in either HEK293T or HCT116 cells. Briefly, as part of our BioPlex project (Huttlin *et al*, 2021), we ectopically expressed PINK1-HA and performed interaction proteomics. While we identified the known protein kinase chaperone HSP90 subunits in association with PINK1, we did not detect FBXO7 (**Figure EV 3A, left panel; Table EV1**). Likewise, FBXO7 associated reciprocally with members of the CUL1-SKP1-RBX1, the COP9/Signalosome, and both core particle and regulatory subunits of the proteasome in HEK293T and/or HCT116 cells, as expected (Bader *et al*, 2011; Liu *et al*, 2019; Vingill *et al*, 2016), but neither PINK or Parkin were detected (**Figure EV 2B; Table EV2**) (Huttlin *et al*., 2021) (see **Materials and Methods**). These results are consistent with our previous Parkin interaction proteomics experiments that failed to demonstrate an interaction with FBXO7 during mitophagy induction (Sarraf *et al*., 2013). Among FBXO7 interacting proteins is PSMF1 (also called PI31). We found that purified SKP1-FBXO7 complexes (with or without CUL1 N-terminal domain (NTD) can directly associate with PSMF1, as examined using size exclusion chromatography and analysis of fractions by SDS-PAGE (**Figure EV 3D,E**). To our knowledge, this is the first demonstration that FBXO7 assembled with SKP1/CUL1 can bind directly to PSMF1.

### Mitophagic flux in FBXO7^-/-^ cells

A significant advance has been the development of mitophagy reporters, which cumulatively measure flux of mitochondrial turnover through to the final stages of mitophagy: the fusion of mitochondria-laden autophagosomes with lysosomes (Lazarou *et al*., 2015). mKeima is a fluorescent protein that undergoes a pH-dependent Stokes shift, and delivery of a Keima-tagged cargo from the cytoplasm to the acidic interior of the lysosome can be monitored by measuring the ratio of emission at 620 nm with maximal excitation at 440 nm or 586 nm under neutral or acidic conditions, respectively, by flow cytometry or microscopy (**Figure 3A**). To monitor differences in mitophagic flux upon deletion of FBXO7, we employed mtKeima, an mKeima protein targeted to the mitochondria via a COX8-matrix targeting sequence (Ordureau *et al*., 2020).

**Figure 3.**
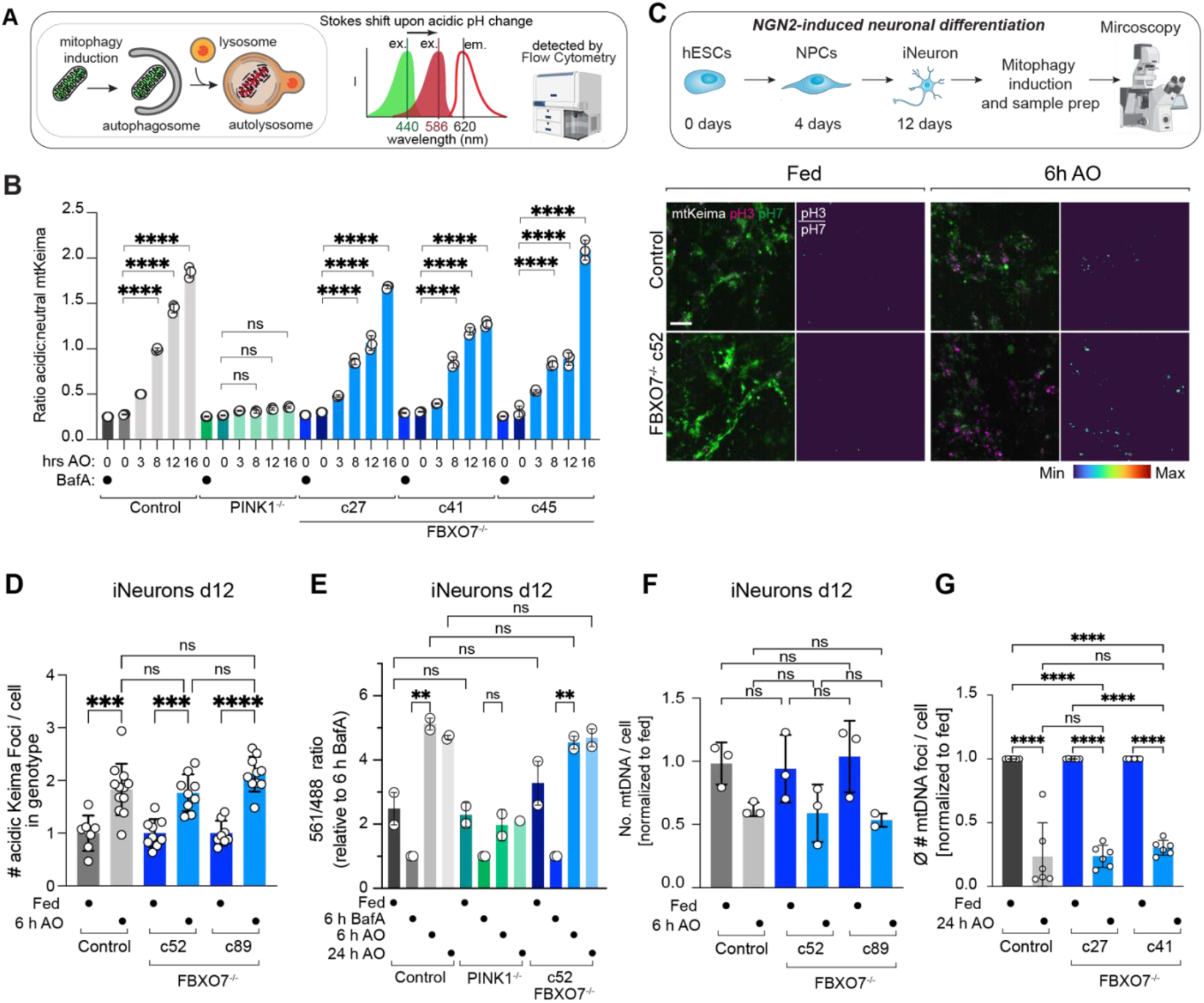
Mitophagic flux in iNeurons and HeLa cells lacking FBXO7. (**A**) Scheme depicting mitophagic flux assay using mtKeima. See text for details. (**B**) Mean Acidic:Neutral mtKeima per-cell ratios measured by flow cytometry for HeLa cells expressing Parkin ndicating the number of hours treated with AO (Antimycin A (5 μM) and Oligomycin (10 μM)) or three hours with 25 nM BafilomycinA (BafA). Error bars depict S.D. from triplicate measurements. Two-way ANOVA with multiple comparisons; p(****)<0.0001. (**C**) Indicated hESCs expressing mtKeima were differentiated to iNeurons (day 12) and were subjected to image analysis. Scale bar, 5 μm. (**D**) The number of red-shifted Keima foci per cell is plotted, where each dot represents the average foci-number per image stack, originating from 7-12 image stacks of iNeuron differentiations. Error bars represent S.D.. One-way ANOVA with multiple comparisons; p(***)=0.0001, p(****)<0.0001. (**E**) mtKeima mitophagy flux readout of indicated iNeuron genotypes. Cells were treated with AO for either 1 or 6h and mitophagic flux measured by flow cytometry (>10,000 cells). Pooled data from 2 biological replicates is shown, normalized to 6h BafA treated cells. Error bars depict S.D.. One-way ANOVA with multiple comparisons; p(**)=0.0011 and 0.0049. (**F**) Bar graph showing the number of mtDNA puncta/cell with or without treatment with AO (6 h) in control or FBXO7^-/-^ day 12 iNeurons. Error bars depict S.D. from triplicate differentiations. One-way ANOVA with multiple comparisons. (**G**) Bar graph showing the number of mtDNA puncta/cell with or without treatment with AO (6 h) in HeLa control or FBXO7^-/-^ cells. Error bars depict S.D. from triplicate replicates. One-way ANOVA with multiple comparisons; p(****)<0.0001.

In Parkin-expressing HeLa cells, the ratio of acidic-Keima (measured with excitation at 561 nm and emission 620 nm) to neutral Keima (ex. 405 nm, em. 603 nm) increases as cells are treated with AO (**Figure 3B, Figure EV 3B**). This mitophagic flux is PINK1-dependent as deletion of PINK1 abolishes this shift (**Figure 3B, Figure EV 3B**). As expected, there is little flux without depolarization with AO, and inhibition of the lysosomal V-ATPase with BafilomycinA1 (BafA) restores the mtKeima acidic:neutral ratio to levels similar to that observed in fed cells (**Figure EV 3B**). In three FBXO7^-/-^ clones, AO-dependent mitophagic flux is similar to control cells with increasing flux observed with increasing length of AO treatment (**Figures 3B; Figure EV 3B**). We validated the mtKeima flux results using the mitochondrial DNA clearance assay (Heo *et al*., 2015; Lazarou *et al*., 2015). Staining for mtDNA foci per cell after 24 h AO treatment in control and FBXO7^-/-^ HeLa cells showed a significant decrease in mtDNA number in all samples (**Figure 3G**). The remaining mtDNA levels were indistinguishable between control and FBXO7^-/-^ cells (**Figure 3G**).

We also measured mtKeima foci and flux in control and FBXO7^-/-^ iNeurons at day 12 of differentiation (**Figure 3C-E**). As with HeLa cells, red-shifted mtKeima foci increased ∼2-fold after 6 h AO treatment in control and FBXO7^-/-^ cells (**Figure 3C,D**). Using flow cytometry, mitophagic flux (normalized to BafA) after 6 and 24 h AO treatment was comparable between control and FBXO7^-/-^ iNeurons, and was absent in PINK1^-/-^ cells (**Figure 3E**). Additionally, we investigated mtDNA turnover in d12 control and two independent FBXO7^-/-^ iNeuron clones. Within 6 or 24 h of depolarization of iNeurons, we observed a reduction in the number of mitochondrial DNA/cell, and the effect was independent of the presence or absence of FBXO7 (**Figure 3F,G**). Taken together, these data indicate that FBXO7 is not universally required for depolarization-dependent mitophagic flux in either the HeLa or iNeuron systems.

### Proteomic analysis of HeLa cells lacking FBXO7 during mitophagy

As an alternative to mtKeima for measuring mitophagic flux, we examined the total proteome of depolarized HeLa cells with the expectation that in control HeLa cells, mitochondrial turnover would result in bulk loss of the mitochondrial proteome in depolarized cells. Thus, control or FBXO7^-/-^ HeLa cells expressing Parkin were depolarized for 16 h in triplicate and total cell extracts subjected to 18-plex Tandem Mass Tagging (TMT)-based proteomics (**Figure 4A, Table EV3,4**) (Li *et al*, 2021). Replicates were highly correlated, with correlation coefficients greater that 0.95 and Principal Component Analysis (PCA) revealed tight clustering of replicates, with the major feature separating the samples being AO-treatment (**Appendix Figure S3A,B**). Through hierarchical clustering (**Figure EV 3C**), we identified major alterations in the abundance of mitochondria in response to depolarization (**Figure 4B, Appendix Figure S3D-I**). Indeed, the majority of proteins annotated as mitochondrial were found to have reduced levels in control cells in response to depolarization, as indicated in the overlap of mitochondrial proteins on the volcano plots for the ∼8000 proteins quantified (leftward skew of red dots in **Figure 4B**, left panel). Importantly, the patterns of protein abundance were indistinguishable from control for two independent FBXO7^-/-^ cell lines, with leftward skew of mitochondrial proteins in the volcano plot (**Figure 4B,C**). In similar fashion, ∼4.3% of detected mitochondrial proteins were significantly more abundant after AO treatment across all genotypes, compared to ∼67.9% of significantly decreased proteins. The plurality (49%) of proteins with significantly decreased abundance was shared between both knockout clones and the KO, supporting a robust mitophagy response even in the absence of FBXO7 (**Appendix Figure S3J,K**). When calculating the difference (delta = Δ log_2_ FC [16h AO / Fed]) between FBXO7^-/-^ and Control for each detected protein and compare the ranked abundance of mitochondrial vs all other proteins, mitochondrially localized proteins are evenly distributed across the whole range, arguing that the proteome of both Control and FBXO7^-/-^ cells remodel in similar fashion after mitophagy treatment (**Appendix Figure S3L**, green dots). Proteins with altered abundance were enriched for mitochondrial matrix, inner membrane, outer membrane and oxidative phosphorylation categories, consistent with autophagy of entire organelles (**Figure 4C**). The behaviour of other organelles and large protein complexes were similar for both control and FBXO7^-/-^ cells (**Figure 4C**). Moreover, both FBXO7 clones displayed similar behaviour to each other and to the control, as indicated by correlation analysis (Pearson’s R=0.82, 0.86 and 0.74, respectively) (**Figure 4D,E**). These results are consistent with the absence of a discernible alteration in depolarization-dependent mitophagic flux in cells lacking FBXO7. A previous report proposed that depletion of FBXO7 resulted in accumulation of PINK1 proteins levels under basal conditions (Huang *et al*., 2020), but in our experiments, PINK1 levels in the two FBXO7^-/-^ clones were unaffected based on TMT intensities (**Figure 4F**).

**Figure 4.**
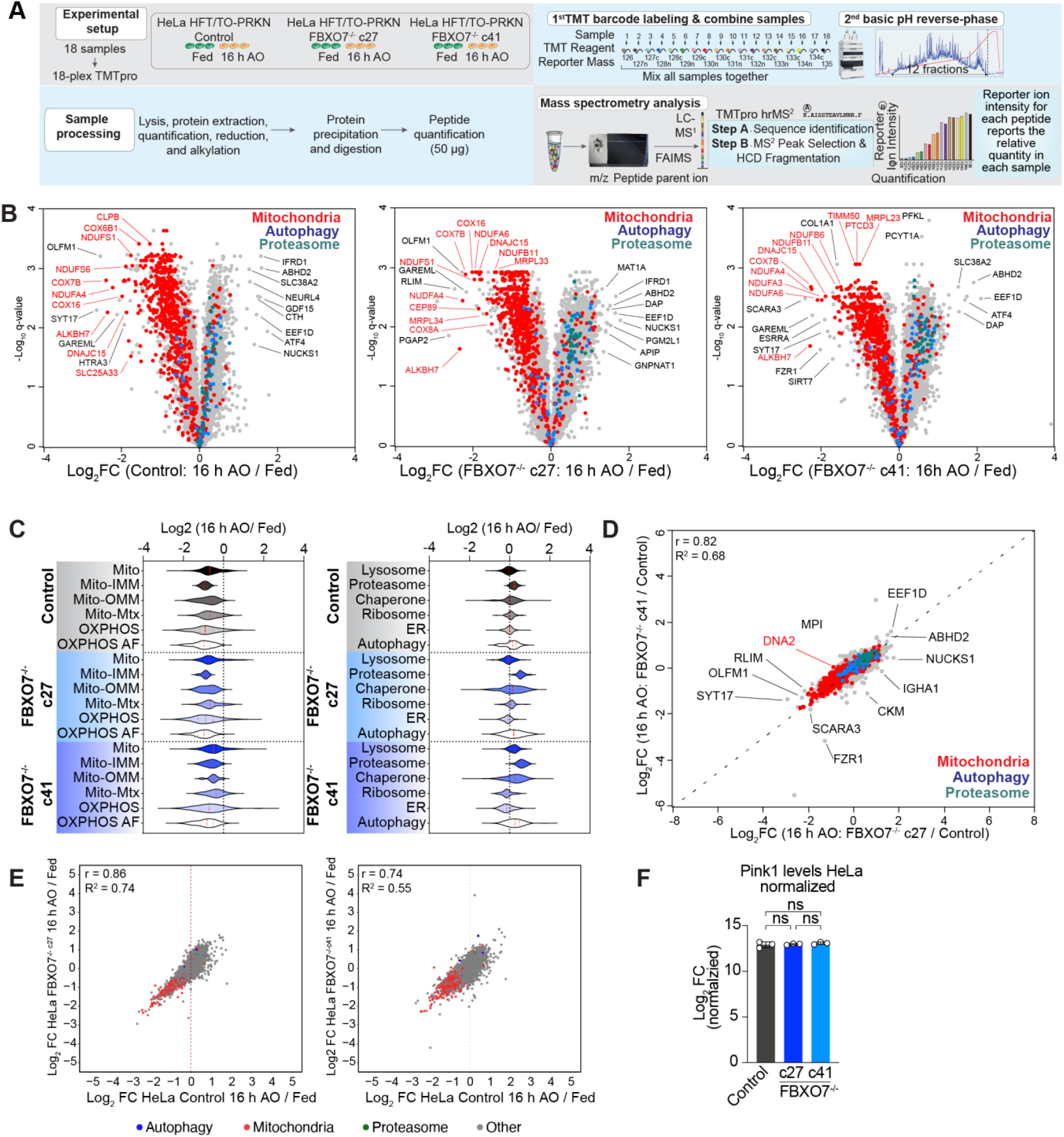
Proteomic analysis of HeLa cells lacking FBXO7 during mitophagy. (**A**) Workflow for analysis of total protein abundance in HeLa cells expressing Parkin with and without depolarization with AO (16 h). Cell extracts were digested with trypsin prior to 18-plex TMT labelling and analysis by mass spectrometry. (**B**) Volcano plots [Log_2_ FC (16 h AO / Fed) versus -Log_10_(q-value)] for control or FBXO7^-/-^ cell lines with or without treated with AO (16 h). Red dots, mitochondrial proteins; blue dots, autophagy proteins; green dots, proteasome proteins; grey dots, remainder of the proteome. (**C**) Violin plots [Log_2_ (16 h AO/Fed)] of control or FBXO7^-/-^ cells depicting alterations in the abundance of mitochondrial protein (left panel) or specific organelles or protein complexes (right panel). (**D**) Correlation plot of c27 and c41 FBXO7^-/-^ clones. Log_2_FC (16 h AO for each clone relative to control cells) is plotted. (**E**) Correlation plots [Log_2_ FC (16 h AO / Fed)] of the proteome of FBXO7^-/-^c27 or c41 clones against control. Red dots, mitochondrial proteins; blue dots, autophagy proteins; grey dots, remainder of the proteome. (**F**) PINK1 levels in control cells and in two FBXO7^-/-^ clones were measured by TMT-proteomics in fed cells (n=3).

### Proteomic analysis of human ES cells lacking FBXO7 during neurogenesis

As an initial unbiased approach to examining FBXO7 function in iNeurons, we performed 18-plex TMT proteomics on control and two clones of FBXO7^-/-^ cells at day 0, 4 and d12 of differentiation (**Figure 5A, Table EV5-8**). We have previously demonstrated dramatic remodelling of mitochondria around day 4 of differentiation to support a switch from glycolysis to oxidative phosphorylation, which is accompanied by BNIP3L-dependent mitophagy of a fraction of mitochondria (Ordureau *et al*, 2021). From the ∼6000 proteins quantified across all replicates and conditions, we found strong correspondence of control and FBXO7^-/-^ cells, with PC1 being driven by differentiation (**Appendix Figure S4A,B**). FBXO7 itself was expressed at comparable levels across the time course but was not detected in FBXO7^-/-^ iNeurons, as expected (**Appendix Figure S4C**). Remarkably, the abundance of the proteome was largely immune to the loss of FBXO7 at both day 4 and day 12 of differentiation (**Figure 5B-D, Appendix Figure S4D**). Moreover, the abundance of mitochondrial proteins (including inner and out membrane and OXPHOS components) were unchanged relative to control cells at day 4 or 12 of differentiation (**Figure 5B, Appendix Figure S4D**). Correlation analysis of Control and FBXO7-/-cells [iNeuron d12 / hESC d0] did show high similarity between the proteomes, as well as for subsets of mtUPR or autophagy markers (**Appendix Figure S4E,F**).

**Figure 5.**
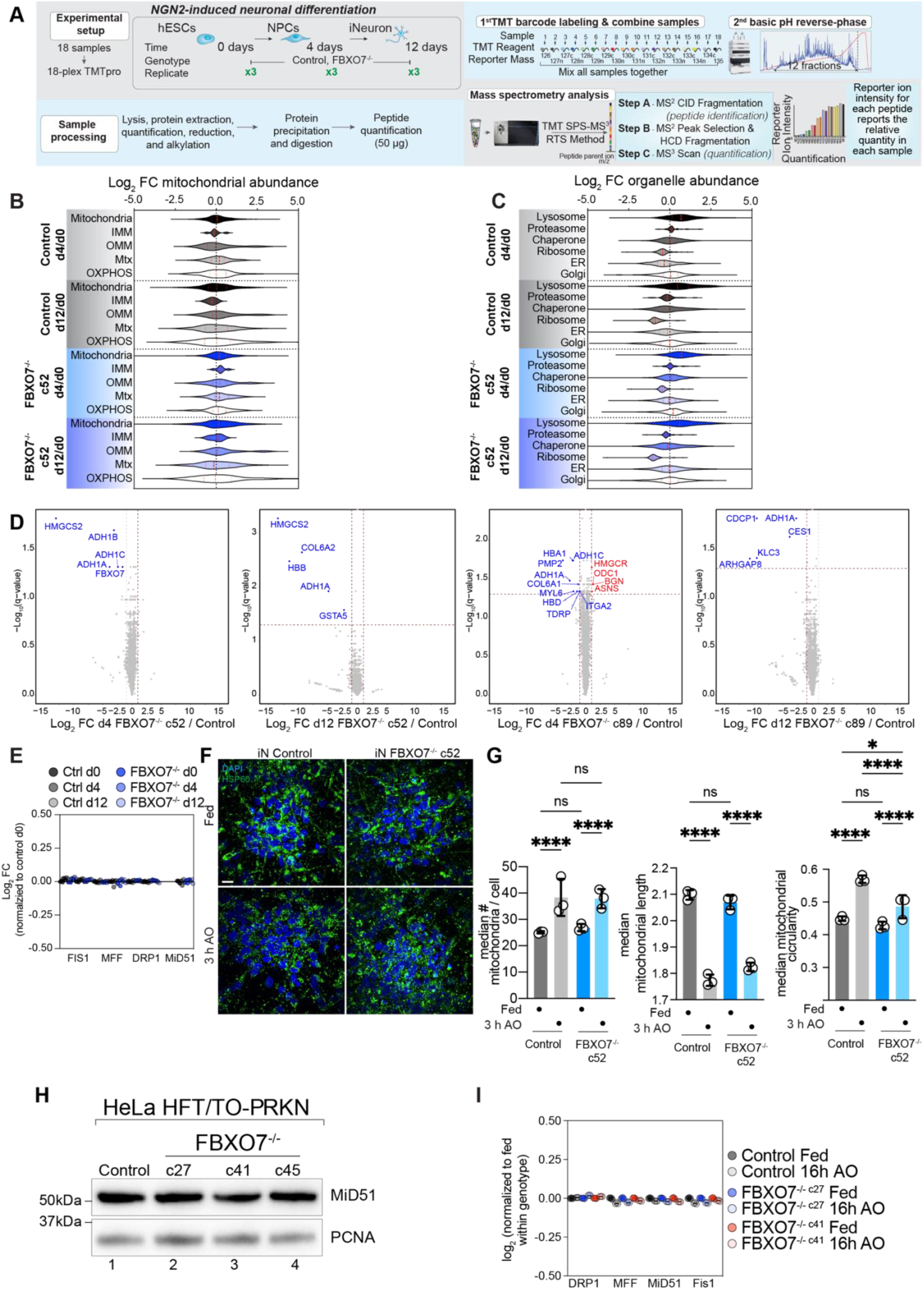
Proteomic analysis of human ES cells lacking FBXO7 during neurogenesis. (**A**) Workflow for analysis of total protein abundance in ES cells undergoing NGN2-driven neurogenesis with or without FBXO7. Cell extracts were digested with trypsin prior to 18-plex TMT labelling and analysis by mass spectrometry. (**B**) Log_2_ FC for the indicated mitochondrial protein-groups in control or FBXO7^-/-^ cells at day 0, 4 or 12 during neurogenesis is shown in violin plots. (**C**) Log_2_ FC for the indicated cellular organelle proteins in control or FBXO7^-/-^ cells at day 0, 4 or 12 during neurogenesis is shown in violin plots. (**D**) Volcano plots [Log_2_ FC (FBXO7^-/-^/Control) versus -Log_10_(q-value)] for FBXO7^-/-^ (c52 and c89) and control cells at day 4 (left panel) or day 12 (right panel) of differentiation. Proteins showing decreased or in creased abundance are shown as blue or red dots. (**E**) Relative abundance of DNM1L (Drp1), MFF, and FIS1 in control or FBXO7^-/-^ cells at day 0, 4 or 12 during neurogenesis. (**F,G**) Mitochondrial morphology in iNeurons (iN) was assessed using confocal imaging after staining cells with HSP60 to detect mitochondria and DAPI to identify nuclei, in either fed cells or cells treated with AO (3 h) (left panel). The number of median # of mitochondria/cell, median mitochondrial length, and median mitochondrial circularity is shown. Error bars depict S.D. from biological triplicate experiments (9-12 image stacks each), as shown in panel I left, center, and right, respectively. Two-way ANOVA with multiple comparisons; p(*)=0.0416, p(****)<0.0001(**H**) Western Blot analysis on HeLa control and FBXO7^-/-^ whole cell lysate, probed for the mitochondrial fission adapter MiD51. (**I**) Relative abundance of DNM1L, MFF, MiD51 and FIS1 in WT (Control) or FBXO7^-/-^ HeLa cells either in the fed state or after 16 h AO as determined from the proteomics data in **Figure 5A,B**.

Recent reports have directly linked L250P mutations in the dimerization domain of FBXO7 to MiD49/MiD51, a core adapter protein involved in mitochondrial fission (Al Rawi *et al*., 2022). Indeed, DNM1L-mediated mitochondrial fission has been previously indicated in the efficient scission of damaged parts of mitochondria and protection of the remaining healthy mitochondrial network (Burman *et al*, 2017). Cells expressing FBXO7^L250P^ were found to have reduced levels of proteasome subunits as well (Al Rawi *et al*., 2022). Our proteomic data from iNeurons indicates no alterations in the abundance of proteasome subunits or mitochondrial fission and fusion machinery (DNM1L, MFF, and MiD51) in iNeurons lacking FBXO7 (**Figure 5C,E**). To test if loss of FBXO7 causes in morphological changes in the mitochondrial network, we stained day 12 iNeurons with for the matrix marker HSP60 prior to image analysis (**Figure 5F,G**). However, we observed no significant alterations in the number of mitochondria/cell or the mean mitochondrial length in control or FBXO7^-/-^ iNeurons. The mean mitochondrial circularity was slightly reduced in FBXO7^-/-^ iNeurons treated with AO for 3h (**Figure 5F**). Analogous analyses of mitochondria and fission/fusion proteins also did not reveal significant alterations in response to loss of FBXO7 in two independent clones of HeLa cells (**Figure 5H,I**). However, there was a ∼10% decrease in the abundance of proteasome subunits in HeLa cells lacking FBXO7 in the absence of depolarization, but an increase after mitophagy induction (**Appendix Figure S3E, Figure 4C**). If this response is a compensatory mechanism of the knockout cell line to cope with cytotoxic insults is an intriguing idea. Last, we examined whether FBXO7^-/-^ is required for the maintenance of the mitochondrial organelle pool during differentiation. Previous work has shown significant metabolic rewiring during the differentiation from hESC to iNeurons, including a switch from glycolysis to oxidative phosphorylation as the cells main energy source (Ordureau *et al*., 2021). Since this switch is accompanied by increased mitophagy, we set out to see if the absence of FBXO7 would lead to accumulation of mitochondria and/or impair differentiation. FBXO7^-/-^ cells did not display obvious differences in the abundance of mitochondrial proteins, when normalized to day 0 (**Figure 5B, Appendix Figure S4D**). Likewise, abundance of other organelles was unchanged (**Figure 5C**). Finally, neither differentiation markers or autophagy proteins displayed obvious changes in control versus the two FBXO7^-/-^ clones as measured by TMT-proteomics and displayed using a T^2^-statstic (Ordureau *et al*., 2021) (**Appendix Figure S5A-F**).

### FBXO7^-/-^ are susceptible to ^mt^UPR

Last, we investigated the role of FBXO7 in response to the mitochondrial unfolding response (^mt^UPR) via treatment of the mitochondrial HSP90 inhibitor Gamitrinib-triphenylphosphonium (G-TPP), which is also known to induce protein misfolding in the matrix and activate the PINK1/Parkin pathway (Fiesel *et al*, 2017; Kataura *et al*, 2023; Munch & Harper, 2016). We treated HeLa or HEK293T Control and FBXO7^-/-^ cells expressing Parkin with 10μM G-TPP (8 or 12h) and examined pUb accumulation using immunoblot (**Figure EV 4A,B**). Cells lacking FBXO7 still responded to depolarization by activation of pUb accumulation, indicating that FBXO7 is dispensable for PINK1/Parkin activation in response to ^mt^UPR. Interestingly, loss of FBXO7 did show increased pUb signal in all knockout clones, suggesting that loss of FBXO7 may prime cells for enhances mtUPR signaling to the PINK1/Parkin pathway. As an orthogonal approach, GFP-Parking expressing HeLa Control and FBXO7^-/-^ were treated with G-TPP, and image analysis used to examine the co-incidence of pUb and GFP-Parkin with mitochondria (**Figure EV 4C**). The average number of pUb after G-TPP was ∼ 4-5 times lower compared to AO treated cells, in line with the lower acute mitotoxic nature of the drug (**Figure EV 4D**). However we observed a ∼10 % increase in pUb foci in FBXO7^-/-^ cells expressing GFP-Parkin when normalized to all Parkin expressing cells (**Figure EV 4E**).

Next, we tested if Parkin is required for the FBXO7-mediated changes in pUb recruitment after G-TPP treatment. HeLa HFT/TO-PARK2 Control and FBXO7^-/-^ cells were treated with G-TPP without and with the addition of doxycycline, thus controlling for Parkin expression. Immunofluorescence staining for pUb revealed the absence of signal in cells lacking Parkin [(–) dox samples], while Parkin-expressing cells responded to HSP90 inhibition with pUb foci appearing after 8h of treatment (**Figure EV 4F**). Analysis of pUb intensity over G-TPP treatment-timecourse suggested potential small differences in pUb kinetics, especially during early states of G-TPP-mediated stress (4-6 h of treatment), albeit variability between FBXO7^-/-^ clones was observed (**Figure EV 4G,H**). After 8h of G-TPP, the normalized ratio of pUb/DNA intensities were on average 20% higher across the knockouts compared to control cells, consistent with immunoblotting.

## Discussion

Despite strong genetic evidence linking mutations in FBXO7 with parkinsonian-pyramidal syndrome (Di Fonzo *et al*., 2009; Houlden & Singleton, 2012; Paisan-Ruiz *et al*., 2010), our understanding of the cellular roles of FBXO7 are limited. An early study reported biochemical links between FBXO7 and PINK1/Parkin-dependent mitophagy in fibroblasts and SH-SY5Y cells, including interaction of overexpressed FBXO7 with both PINK1 and Parkin (Burchell *et al*., 2013). Using siRNA to deplete FBXO7, it was also concluded that FBXO7 promotes Parkin’s ability to ubiquitylate the outer mitochondrial membrane substrate MFN1 and also promotes clearance of mitochondria by autophagy. These biochemical and physiological results led to the conclusion that FBXO7 functions as a biochemical amplifier of PINK1/Parkin-dependent mitophagy (Burchell *et al*., 2013). However, follow-up studies have been limited, and the field has experienced dramatic advances in the understanding of the biochemical roles of Parkin and PINK1 in promoting mitochondrial clearance by autophagy, thereby providing the tools to examine potential roles of FBXO7 in the pathway.

With these tools, we set out to define where in the pathway FBXO7 might operate to promote mitophagy. However, in both the HeLa cells with overexpressed Parkin and iNeurons with a fully endogenous PINK1/Parkin pathway, we failed to validate a role for FBXO7 in any of several steps in the pathway, including pUb accumulation, Parkin recruitment to the mitochondrial outer membrane, recruitment of autophagy machinery, mitophagic flux as measured by mtKeima, and mitochondrial proteome degradation. The finding that deletion of FBXO7 has no obvious effect on the pathway with either endogenous or overexpressed Parkin in iNeurons and HeLa cells indicates that FBXO7 does not play a general or universally required role in the pathway. A previous study largely based on overexpression concluded that FBXO7 may bind and regulate PINK1 levels, with <2-fold increases in PINK1 by immunoblotting in FBXO7-depleted cells (Huang *et al*., 2020). However, our quantitative proteomics experiments (**Figure 4**) indicated that deletion of FBXO7 has no effect on PINK1 abundance, consistent with the finding that pUb accumulation in response to depolarization is unaffected in FBXO7^-/-^ cells. Moreover, our previous interaction proteomics experiments failed to identify either PINK1 in association with overexpressed FBXO7 (Huttlin *et al*., 2021) or FBXO7 in association with overexpressed Parkin (Sarraf *et al*., 2013). Thus, our functional and biochemical results do not support a biochemical linkage between FBXO7 and the PINK1/Parkin pathway.

In order to broadly examine FBXO7^-/-^ iNeurons for pathways that are affected, we performed global proteomic analysis during neurogenesis *in vitro*. However, we did not observe any alterations in pathways linked with mitochondria, or other quality control pathways, although a reduction in the abundance of alcohol dehydrogenase enzymes among a small number of other proteins, was observed. Whether these signatures are related to FBXO7 function remains to be examined. While a recent study linked the L250P mutation in the dimerization domain of FBXO7 with alterations in proteins associated with mitochondrial fusion and fission (Al Rawi *et al*., 2022), we did not observe alterations in either the abundance of these proteins (MiD51, MFF, FIS1) nor did we observe changes in mitochondrial morphology in either HeLa cells or iNeurons lacking FBXO7. However, based on the G-TPP results, we cannot exclude the possibility that stress response pathways distinct from those examined here, and possibly in specific cell types, may employ FBXO7 to promote mitochondrial health.

An apparent ortholog of FBXO7 was identified in *Drosophila* – referred to as *nutcracker, ntc* – and shown to associate with components of the core particle of the proteasome, as well as with DmPI31 (Bader *et al*., 2011). Association of human FBXO7 with proteasome components and PSMF1/PI31 have also been observed in both focused studies and in our previous interaction proteomic studies (Al Rawi *et al*., 2022; Huttlin *et al*., 2021; Vingill *et al*., 2016). In this context, PI31/DmPI31 and FBXO7/*ntc* have been proposed to functionally link proteasome trafficking via dynein motors in neuronal processes (Liu *et al*., 2019). Of note, in our HeLa proteomics, proteasome proteins appeared to be overrepresented in FBXO7^-/-^ proteomes after mitophagy treatment (**Figure EV4I**). Our interaction proteomics analysis has also confirmed association of FBXO7 with PSMF1, CRL, COP9/Signalosome, and proteasome core subunits, and demonstrated that association of FBXO7-SKP1-CUL1^NTD^ with PSMF1 is a direct interaction (**Figure EV 3D,E**). Further studies are required to elucidate the relationship between FBXO7 and parkinsonian-pyramidal syndrome, to understand any functional relationships between FBXO7 and other Parkinson’s disease risk alleles, and to examine whether FBXO7’s physical association with the proteasome is linked with disease.

## Acknowledgments

This work was supported by Aligning Science Across Parkinson’s (ASAP) (J.W.H. and B.A.S.), the NIH (R01 NS083524 to J.W.H. and RO1GM132129 to J.A.P.), The Max Planck Society (B.A.S.), and the Harvard Medical School Cell Biology Initiative for Molecular Trafficking and Neurodegeneration. Michael J Fox Foundation administers the grant ASAP-000282 on behalf of ASAP and itself. For the purpose of open access, the author has applied a CC-BY public copyright license to the Author Accepted Manuscript (AAM) version arising from this submission. We thank Laura Pontano Vaites for help with the BioPlex 3.0 interactome analysis. We thank Jennifer Waters, Talley Lambert, Federico Gasparoli and Rylie Walsh in the Nikon Imaging Center and the Cell Biology Microscopy Facility (CBMF) at Harvard Medical School for microscopy support and technical support.

## Data Availability

Proteomic data and analysis files part of this study are deposited at ProteomeXchange Consortium by the PRIDE partner (Deutsch *et al*, 2020; Perez-Riverol *et al*, 2022). The PRIDE project identification number is PXD037797 and can be accessed for reviewers under username; password 65wg8SiI. Macros and pipelines used in this work can be found on GitHub (https://github.com/harperlaboratory/FBXO7.git) or Zenodo (https://zenodo.org/record/7803294). Raw files associated with this work are deposited on Zenodo (https://zenodo.org/record/7803294).

## Author Contributions

Study design: J.W.H., F.K., E.A.G.; Data collection: F.K., E.A.G., J.A.P., J.C.P., J.Z., Y.Z., F.A.; Data recording, analysis and interpretation: F.K., E.G., I.R.S., J.A.P., J.Z., Y.Z., F.A., B.A.S.; Manuscript preparation: J.W.H., F.K. with input from all authors.

## Disclosure and competing interest statement

J.W.H. is a consultant and founder of Caraway Therapeutics and a founding scientific advisory board member of Interline Therapeutics. No other authors declare a conflict of interest.

**Figure EV 1.**
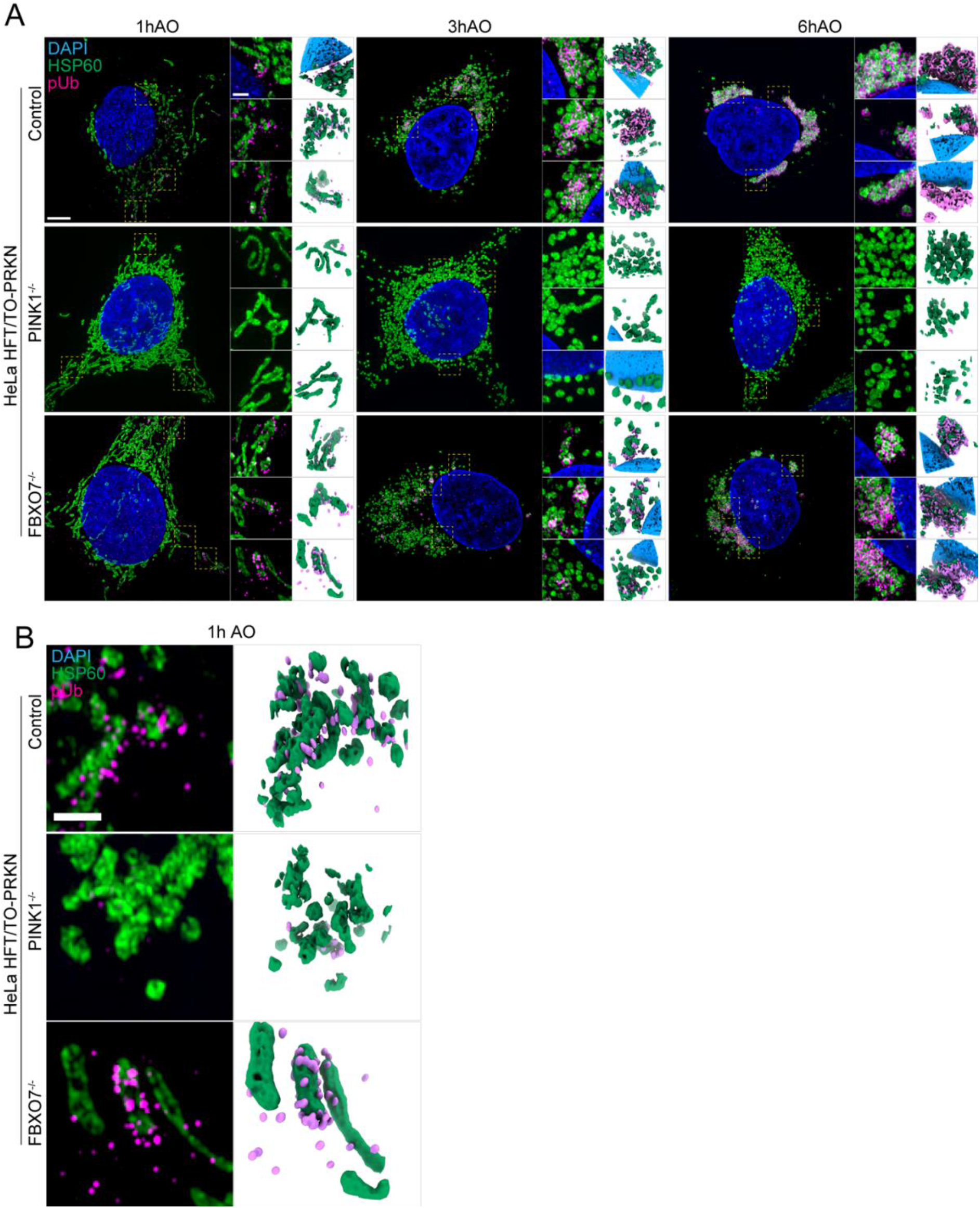
Super-resolution pUb detection in HeLa cells in response to mitochondrial depolarization. 3D-SIM images of HeLa control, PINK1^-/-^ and FBXO7^-/-^ cell lines after 1h, 3h or 6h, AO-induced mitophagy (related to **Figure 2A,B**). Cells were stained for nuclear DNA (DAPI), mitochondria (HSP60) and pUb. (**B**) Zoom-ins of regions of interested after 1h AO-induced mitophagy are depicted and 3D-surface renderings of insets are shown on the right. Scale bar = 5 μm or 1 μm.

**Figure EV 2.**
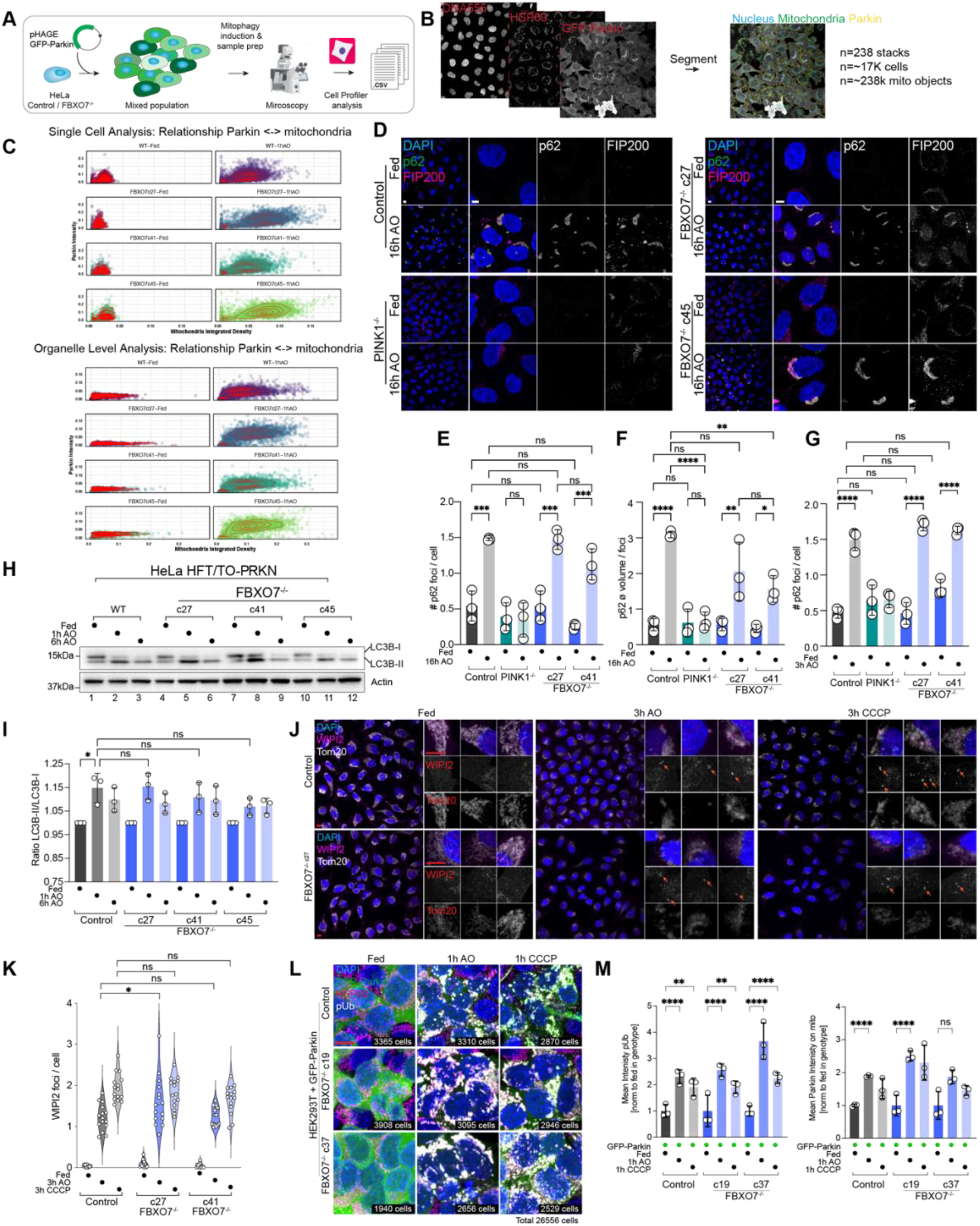
Parkin and autophagy regulator recruitment in FBXO7^-/-^ cells. (**A**) Overview of methods for analysis of Parkin recruitment to damaged mitochondria. (**B**) Cells were imaged for GFP-Parkin, nuclei, and mitochondria and segmented to facilitate analysis of Parkin recruitment to mitochondria in single cells / mitochondrial organelles. (**C**) Std mitochondrial signal vs Parkin intensity and Std Parkin vs mitochondria with or without 1h AO-induced mitophagy are shown. Data from three replicates, including a total of 238 image stacks, containing 16888 cells and 237848 mitochondrial objects. (**D**) Recruitment of p62 to mitochondria (not stained) in control or FBXO7^-/-^ HeLa cells with or without treatment with AO (16 h) was examined by confocal imaging. Scale bar = 10 and 5 μm. (**E,F**) Quantification of cells in panel D. Assays were performed in biological triplicate with 4 image stacks taken per repeat. n = 6712 cells. Error bars depict S.D.. One-way ANOVA with multiple comparisons; p(***)=0.0002, p(*)=0.0355, p(**)=0.0029, p(****)<0.0001. (**G**) Quantification of #p62 foci normalized per cell in control, PINK1^-/-^ and FBXO7^-/-^ cells with or without 3h AO treatment. Assays were performed in biological triplicate with 4 image stacks taken per repeat. n = 12646 cells. Error bars depict S.D.. One-way ANOVA with multiple comparisons; p(****)<0.0001. (**H**) Control or FBXO7^-/-^ HeLa cells expressing Parkin were either left untreated or incubated with AO for 1 or 6 h and extracts subjected to immunoblotting with α-LC3B and α-Actin as a loading control. (**I**) The ratio of LC3B lipidation (LC3B-II/LC3B-I) was quantified in 3 biological triplicate experiments. One-way ANOVA with multiple comparisons. p(*) = 0.0104. (**J**) Immunofluorescence staining of WIPI2 (magenta) and Tom20 (grey) in HeLa Control and FBXO7-/-cells after mitophagy induction. Arrows depict WIPI2 foci on the mitochondrial staining. Scale bar = 10 μm (overview) and 10 μm (insets). (**K**) Evaluation of mitochondrially localized WIPI2 foci per cell based on images shown in **J**. Data based on 15 image stacks from three technical replicates. One-way ANOVA with multiple comparisons. p(*) = 0.0331. (**L**) Immunofluorescence staining of HEK293T Control and FBXO7-/-cells expressing GFP-Parkin for HSP60 (mitochondria, magenta), pUb (grey) and GFP (Parkin, green) after mitophagy induction for indicated times. Scale bar = 10 μm. Number of individual cells analyzed per condition is indicted in the right bottom of each image. (**M**) Analysis of pUb intensity (left) and Parkin intensity (right) in the mitochondrial mask. Error bars depict S.E.M.. 2-way ANOVA with multiple comparisons; p(**)=0.001, p(****)<0.0001.

**Figure EV 3.**
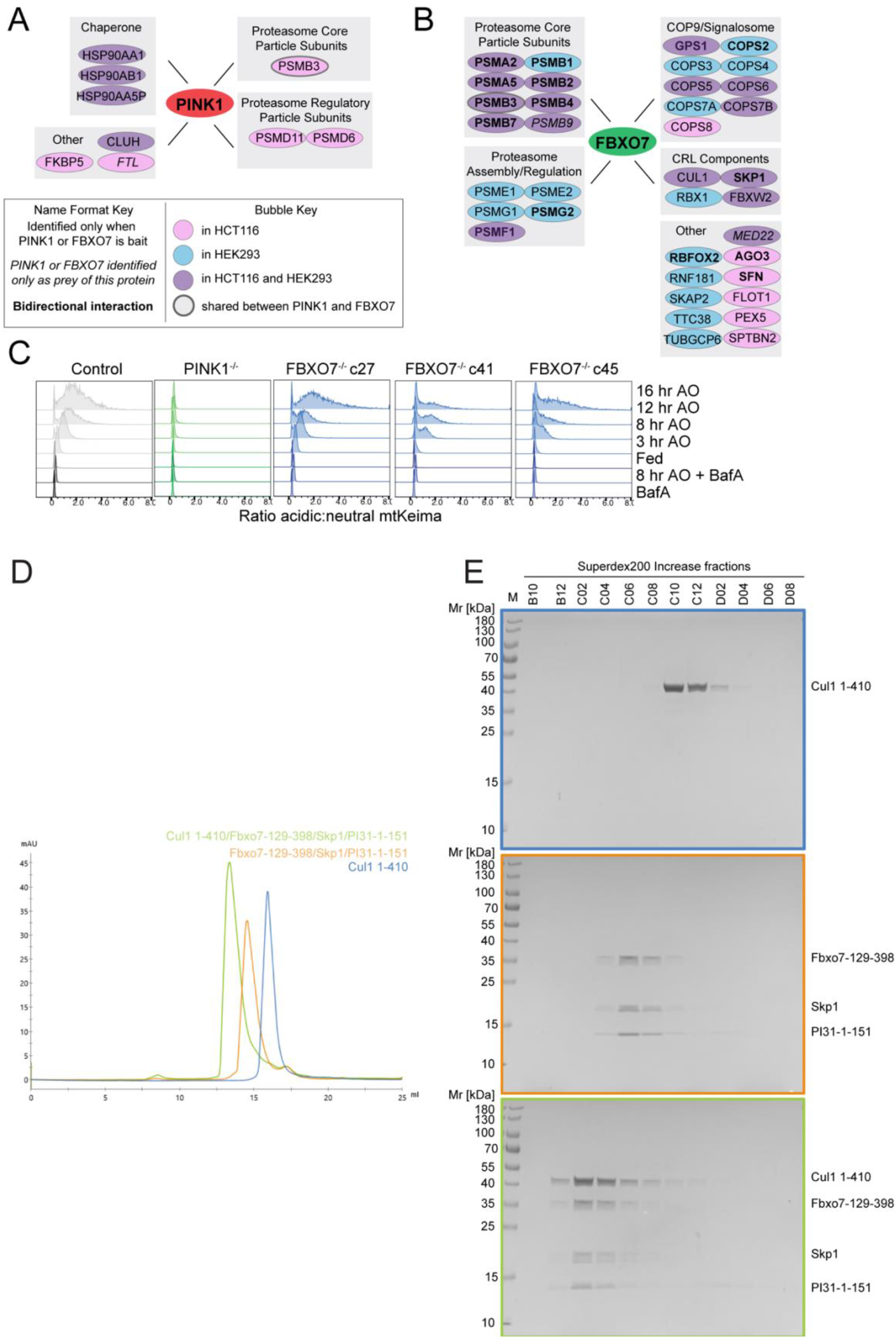
Interaction analysis of FBXO7. (**A**) Interaction network analysis of PINK1 (left) and FBXO7 (right) based on interaction proteomics data from our BioPlex Interactome (Huttlin *et al*., 2021) (see METHODS for details). Figure key can be found beneath figure. (**B**) Summary of major interactions observed for FBXO7 in the context of the relevant protein complexes. (**C**) Ridgeline plots of mtKeima-shift analysis in HeLa control, PINK1^-/-^ and FBXO7^-/-^ cell lines treated with AO at the indicated times. All lines are normalized to the BafA sample. (**D**) Size-exclusion chromatography traces of indicated FBXO7 complexes. (**E**) SDS-PAGE gel analysis of purified SKP1-FBXO7 complexes. Box colours match traces from **D**.

**Figure EV 4.**
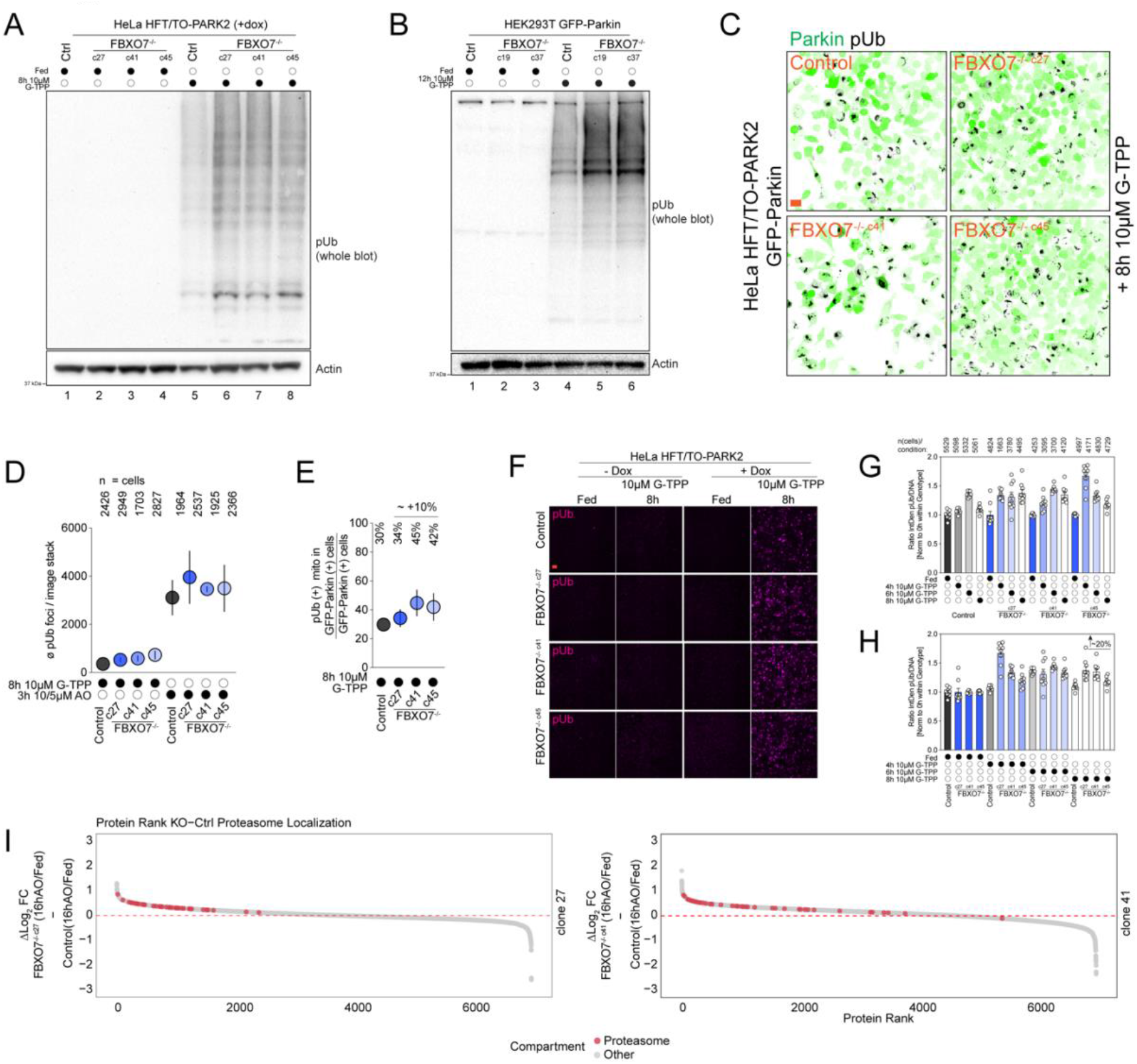
Analysis of FBXO7^-/-^ response to G-TPP. (**A**) Immunoblot for pUb of HeLa Control and FBXO7^-/-^ cells after mitophagy induction using G-TPP for indicated times. (**B**) Immunoblot for pUb of HEK293T Control and FBXO7^-/-^ cells after mitophagy induction using G-TPP for indicated times. (**C**) Immunofluorescence images of HeLa Control and FBXO7^-/-^ cells after G-TPP treatment. Staining for Parkin (GFP, green) and pUb (grey) are shown. Scale bars = 10 μm. (**D,E**) Evaluation of mitophagic response to G-TPP in Hela Control and FBXO7^-/-^ cells, based on images from **C**. ø number of pUb per image stack are plotted. Number of analysed cells for each condition is indicted in **D**. Error bars depict S.E.M.. (**D**) Ratio of pUb-positive mitochondria in GFP-Parkin-positive cells divided by total number of GFP-Parkin-positive cells as proxy for mitophagy induction rate is plotted. Ø % numbers for each genotype are indicted above the plot. Error bars depict S.E.M.. (**F**) Immunofluorescence images of HeLa Control and FBXO7^-/-^ cells after G-TPP timecourse treatment ± doxycycline (= Parkin expression). Staining for pUb (magenta) is shown. Scale bars = 25 μm. (**G,H**) Quantification of pUb / DNA integrated density normalized to 0h treatment within genotype. Panel **H** depicts the measurements sorted by treatment time. N(cells) analysed per condition is indicated at the bottom of the graph. Error bars depict S.E.M.. (**I**) Ranked protein abundance of Δ log2 (KO[16h AO/Fed] – Control [16h AO/Fed]) for FBXO7^-/-^ clone 27 (top) and clone 41 (bottom). Proteasomal annotated proteins are depicted red, all other detected proteins in grey.

## Materials and Methods

All details and catalogue numbers can be found in the **Materials Table (Table EV9)**. Protocols associated with this work can be found on protocols.io (dx.doi.org/10.17504/protocols.io.kxygx99pwg8j/v2)

## Cloning and plasmid generation

The construction of lentiviral expression constructs used in this study have been previously described: pHAGE-mt-mKeima in (Heo *et al*., 2015), PB-mt-mKeimaXL (Ordureau *et al*., 2021) and pHAGE-GFP-Parkin in (Ordureau *et al*., 2018).

## Cell culture, generation of lentiviral stable cell lines

HeLa Flip-In T-Rex (HFT) expressing doxycycline-inducible WT-Parkin (HeLa HFT/TO-PRKN-WT) cells (Heo *et al*., 2015) and HEK293T were maintained in Dulbecco’s modified Eagles medium (DMEM), supplemented with 10% vol/vol fetal bovine serum (FBS), 5% vol/vol penicillin-streptomycin (P/S), 5% vol/vol GlutaMAX and 5% vol/vol Non-essential amino acids (NEAA) at 37°C, 5% O_2_.

Stable cell lines were generated using lentivirus generated from HEK293T cells. pHAGE-mt-mKeima or pHAGE-eGFP-Parkin were co-transfected in HEK293T cells together with the lentiviral vectors pSpax2 and pMD2.1 using Lipofectaime LTX reagent (Thermo Fisher Scientific, 15338100), according to manufacturer’s instructions. Virus was harvested, filtered (0.45 μm) and added 8μg/ml polybrene and target cells infected for 18 h. Antibiotic selection was performed using 1 μg/ml puromycin or FACS sorting and cells verified using immunoblotting or fluorescence microscopy.

If not stated otherwise, Parkin expression in HeLa HFT/TO-PRKN was induced using 2 μg/ml doxycycline o/n, and subsequent mitophagy induced using 5μM Antimycin A/10μM Oligomycin A (==AO), 1μM CCCP or 10μM G-TPP for the indicated times.

H9 ES cells harboring the mitochondrial mt-mKeima flux reporter (Ordureau *et al*., 2020) were generated by electroporation of 1×10^6^ cells with 2.5 μg of pAC150-PiggyBac-matrix-mKeima^XL^ along with 2.5 μg of pCMV-HypBAC-PiggyBac-Helper, as described (Ordureau *et al*., 2020). The cells were selected and maintained in E8 medium supplemented with 50 μg/ml Hygromycin and Hygromycin was kept in the medium during differentiation to iNeurons. Mitophagy was induced with either 0.5 or 5μM Antimycin A or 0.5 or 10μM Oligomycin A.

## Gene-Editing and iNeuron differentiation

Generation of HeLa HFT cells lacking FBXO7^-/-^ was facilitated using CRISPR/Cas9 with target sites determined using CHOPCHOP (Labun *et al*, 2019). Guide RNAs were ligated into the px459 plasmid (Addgene plasmid # 62988) and cells transfected using Lipofectaime LTX reagent (Thermo Fisher Scientific, 15338100), according to manufacturer’s instructions. Two days post-transfection, single GFP positive cells were sorted into 96-well dishes containing 300μl full growth medium (composition as described above). Single cells were allowed to grow into colonies, then duplicated for MiSeq analysis and maintenance. Knockout candidates were confirmed by Western blot on whole cell lysates. The sgRNAs were generated using *GeneArt Precision gRNA Synthesis Kit* (Thermo Fisher Scientific) according to the manufacturer’s instruction and purified using RNeasy Mini Kit (Qiagen). The sgRNA target sequence: ACCGATTCACTACAGAGCAT.

The PINK1^-/-^ H9 cells used here, wherein sequences in exon 1 were deleted using CRISPR-Cas9 to create a null allele, were reported previously (Ordureau *et al*., 2018). To generate FBXO7^-/-^ H9 ES or HeLa cells, 0.6 μg sgRNA was incubated with 3 μg SpCas9 protein for 10 minutes at room temperature and electroporated into 2×10^5^ WT H9 cells using Neon transfection system (Thermo Fisher Scientific). Mutants were identified by Illumina MiSeq and further confirmed by Western blotting. For introduction of TRE3G-NGN2 into the AAVS1 site, a donor plasmid pAAVS1-TRE3G-NGN2 was generated by replacing the EGFP sequence with N-terminal flag-tagged human NGN2 cDNA sequence in plasmid pAAVS1-TRE3G-EGFP (Addgene plasmid # 52343). Five micrograms of pAAVS1-TRE3G-NGN2, 2.5 μg hCas9 (Addgene plasmid # 41815), and 2.5 μg gRNA_AAVS1-T2 (Addgene plasmid # 41818) were electroporated into 1×10^6^ H9 cells. The cells were treated with 0.25 μg/ml Puromycin for 7 days and surviving colonies were expanded and subjected to genotyping.

Human ES cells (H9, WiCell Institute) were cultured in E8 medium (Chen *et al*, 2011) on Matrigel-coated tissue culture plates with daily medium change. Cells were passaged every 4-5 days with 0.5 mM EDTA in 1× DPBS (Thermo Fisher Scientific). SpCas9 and AsCas12a/AsCpf1 expression plasmids pET-NLS-Cas9-6xHis (Addgene plasmid # 62934) and modified pDEST-his-AsCpf1-EC, generated by deleting the MBP sequence from plasmid pDEST-hisMBP-AsCpf1-EC (Addgene plasmid # 79007), were transformed into *Rosetta™(DE3)pLysS* Competent Cells (Novagen), respectively, for expression. SpCas9 and AsCas12a/AsCpf1 proteins were purified as described elsewhere (Hur *et al*, 2016; Zuris *et al*, 2015). Briefly, cells expressing SpCas9 [0.5 mM isopropylthio-β-galactoside, 14-hour induction] were lysed in FastBreak buffer (Promega, Inc) and the NaCl concentration adjusted to 500 mM. Extracts were centrifuged at 38,000g for 10 min at 4°C and the supernatant incubated with Ni-NTA resin for 1 hour. The resin was washed extensively with 50 mM Tris pH8.0, 500 mM NaCl, 10% glycerol, 20 mM imidazole, and 2 mM TCEP prior to elution with this buffer supplemented with 400 mM imidazole. Proteins were diluted two volumes/volume in PBS and fractionated on a Heparin-Sepharose column using a 0.1 to 1.0 M NaCl gradient. Cas9-containing fractions were stored in PBS, 20% glycerol, 2 mM TCEP at -80°C. AsCpf1 expression was induced similarly, and cells pelleted by centrifugation. Cells were lysed by sonication in 50 mM HEPES pH7, 200 mM NaCl, 5 mM MgCl_2_, 1 mM DTT, and 10 mM imidazole supplemented with lysozyme (1 mg/ml) and protease inhibitors (Roche complete, EDTA-free). After centrifugation (16,000g for 30 min), the supernatant was incubated with Ni-NTA resin, the resin washed with 2M NaCl, and bound proteins eluted with 250 mM imidazole, and buffer exchanged into lysis buffer lacking MgCl_2_ and imidazole prior to storage at -80°C.

For human ES cell conversion to iNeurons, cells were expanded and plated at 2×10^4^/cm^2^ on Matrigel-coated tissue plates in DMEM/F12 supplemented with 1x N2, 1x NEAA (Thermo Fisher Scientific), human Brain-derived neurotrophic factor (BDNF, 10 ng/ml, PeproTech), human Neurotrophin-3 (NT-3, 10 ng/ml, PeproTech), mouse laminin (0.2 μg/ml, Cultrex), Y-27632 (10 μM, PeproTech) and Doxycycline (2 μg/ml, Alfa Aesar) on Day 0. On Day 1, Y-27632 was withdrawn. On Day 2, medium was replaced with Neurobasal medium supplemented with 1x B27 and 1x Glutamax (Thermo Fisher Scientific) containing BDNF, NT-3 and 1 μg/ml Doxycycline. Starting on Day 4, half of the medium was replaced every other day thereafter. On Day 7, the cells were treated with Accutase (Thermo Fisher Scientific) and plated at 3-4×10^4^/cm^2^ on Matrigel-coated tissue plates. Doxycycline was withdrawn on Day 10.

## Immunoblotting

At the indicated times, ES cells, iNeurons or HeLa cells were washed on ice in 1x PBS, harvested and pellet wash with 1x PBS and resuspended in 8 M urea buffer (8 M urea, 150 mM TRIS pH 7.4, 50 mM NaCl, PhosSTOP phosphatase inhibitor cocktail). Resuspended cell lysates were sonicated for 10 seconds and debris pelleted at 13,000 rpm for 10 min. Protein concentration was determined by BCA assay according to manufacturer’s instructions (Thermo Fisher Scientific, 23227). Indicated amounts of proteins were resuspended in 1xLDS + 100 mM DTT and boiled for 10 minutes at 85°C. Equal amount of protein and volume were loaded run on 4%-20% Bis-Tris, 8% Tris NuPAGE gels for 5 minutes at 100V, 5 min at 150 V and then run at 200 V for the required time. Gels were transferred via wet transfer system onto PDVF membranes for immunoblotting. Chemiluminescence and colorimetric images were acquired using a BioRad ChemiDoc MP imaging system. Images from Western Blots were exported and analysed using Image Lab and ImageJ/FiJi (Schindelin *et al*, 2012).

## Expression and purification, and analysis of SKP1-FBXO7 complexes

All constructs were prepared utilizing standard molecular biological techniques and verified by Sanger sequencing. The cDNAs coding for HsFbxo7-129-398 and Skp1 preceding a second RBS were cloned into pGEX4T1 as previously described for other substrate receptor/Skp1 complexes (Schulman et al., 2000) (pGEX4T1-TEV-HsFbxo7-129-398/HsSkp1). The cDNA coding for HsPI31-1-151 (with N-terminal TEV cleavable His8-tag) was cloned into pRSF1b (pRSF1b-His8-TEV-HsPI31-1-151). For co-expression of Fbxo7/Skp1/PI31 complexes both plasmids were co-transformed into *E. coli* BL21 Rosetta (DE3). Cultures were grown in Terrific Broth medium, and at an OD600 of 0.8 expression was induced with 0.5 mM IPTG, and *E. coli* were further cultured at 18°C for 16 h. Fbxo7/Skp1/PI31 complexes were purified by sequential standard GST and His affinity chromatography. Affinity tags were cleaved by incubation with TEV protease at 4°C for 16h, and the complexes were further purified by preparative size exclusion chromatography (SEC) in buffer A (25 mM HEPES pH7.5 (KOH), 150 mM NaCl, 1 mM DTT) on a Superdex 200 Increase 10/300 GL column. Fractions of interest were pooled, aliquoted, snap frozen in liquid N_2_, and stored at -80°C until further usage. HsCul1-1-410 was expressed as GST-fusion protein and purified as described previously (Hopf et al., 2022).

## Analytical size exclusion chromatography (SEC)

Analytical SEC was carried out on an ÄKTApure system (GE Healthcare) equipped with a Superdex 200 Increase 10/300 GL column (Cytiva), in buffer A (25 mM HEPES pH7.5 (KOH), 150 mM NaCl, 1 mM DTT). In brief, samples (100 μl at a concentration of 45 μM) of HsCul1-1-410, HsFbxo7-129-398/HsSkp1/HsPI31-1-151, or HsFbxo7-129-398/HsSkp1/HsPI31-1-151 + HsCul1-410 (preincubated at 37°C for 10 min before loading) were applied to the column with a flow rate of 1 ml/min, UV absorbance was recorded at 280 nm, fractions of 200 μl were collected, and analyzed by SDS-PAGE.

## Proteomics

### Proteomics–general sample preparation

Sample preparation of proteomic analysis of whole-cell extract from HeLa, hESC, NPC and iNeurons was performed according to previously published studies (Ordureau *et al*., 2021; Ordureau *et al*., 2020). Cells were harvested on ice and plates were washed twice with 1x PBS and detached in 1x PBS using cell scraper. After pelleting at 2000 rpm for 5 min at 4°C, cells were washed 2x with 1x PBS and resuspended in 8 M urea buffer (composition stated above). After sonification for 10 seconds, resuspended cells were pelleted for 10 min at 13000 rpm. Protein concentration was determined using BCA kit (Thermo Fisher Scientific, 23227).

Unless otherwise noted, proteomics and data analysis was performed as described (Ordureau *et al*., 2021; Ordureau *et al*., 2020). Briefly, protein extracts (100 mg) were subjected to disulfide bond reduction with 5 mM TCEP (room temperature, 10 min) and alkylation with 25 mM chloroacetamide (room temperature, 20 min). Methanol–chloroform precipitation was performed prior to protease digestion. In brief, four parts of neat methanol were added to each sample and vortexed, one part chloroform was then added to the sample and vortexed, and finally three parts water was added to the sample and vortexed. The sample was centrifuged at 6,000 rpm for 2 min at room temperature and subsequently washed twice with 100% methanol. Samples were resuspended in 100 mM EPPS pH8.5 containing 0.1% RapiGest and digested at 37 °C for 8h with Trypsin at a 100:1 protein-to-protease ratio. Samples were acidified with 1% Formic Acid for 15 min and subjected to C18 solid-phase extraction (SPE) (Sep-Pak, Waters).

### Proteomics – quantitative proteomics using TMT

Tandem mass tag labeling of each sample (50 mg peptide input) was performed by adding 5 uL of the 25 ng/mL stock of TMTpro reagent along with acetonitrile to achieve a final acetonitrile concentration of approximately 30% (v/v). Following incubation at room temperature for 1 h, the reaction was quenched with hydroxylamine to a final concentration of 0.5% (v/v) for 15 min. The TMTpro-labeled samples were pooled together at a 1:1 ratio. The sample was vacuum centrifuged to near dryness, and following reconstitution in 1% FA, samples were desalted using C18 solid-phase extraction (SPE) (50 mg, Sep-Pak, Waters), according to manufacturer protocol.

Dried TMTpro-labeled peptides (∼300 ug) were resuspended in 10 mM NH_4_HCO_3_ pH 8.0 and fractionated using basic pH reverse phase HPLC (Wang *et al*, 2011). Briefly, samples were offline fractionated over a 90 min run, into 96 fractions by high pH reverse-phase HPLC (Agilent LC1260) through an Aeris peptide xb-c18 column (Phenomenex; 250 mm x 3.6 mm) with mobile phase A containing 5 % acetonitrile and 10 mM NH_4_HCO_3_ in LC-MS grade H_2_O, and mobile phase B containing 90 % acetonitrile and 10 mM NH_4_HCO_3_ in LC-MS grade H_2_O (both pH 8.0). The 96 resulting fractions were then pooled in a non-continuous manner into 24 fractions (as outlined in Figure S5 of (Paulo *et al*, 2016)) and sets of 12 fractions (even or odd numbers) were used for subsequent mass spectrometry analysis. Fractions were vacuum centrifuged to near dryness. Each consolidated fraction was desalted via StageTip, dried again via vacuum centrifugation, and reconstituted in 5 % acetonitrile, 1 % formic acid for LC-MS/MS processing.

TMT proteomics samples were subjected to analysis using a Orbitrap Fusion Lumos Tribrid mass spectrometer (Thermo Fisher Scientific, San Jose, CA) online with Proxeon EASY-nLC1200 liquid chromatography (Thermo Scientific). Peptides were resuspended in 5 % ACN/5 % FA and 10 % of the samples were loaded on a 35 cm analytical column (100mm inner diameter) packed in-house with Accurcore150 resin (150 Å, 2.6 mm, Thermo Fisher Scientific, San Jose, CA) for LC-MS analysis. Peptide separation was performed with a gradient of acetonitrile (ACN, 0.1% FA) from 3-13 % (0-83 min) and 13-28 % (80-83 min) during a 90 min run.

For the HeLa TMTpro proteomic samples, LC-hrMS/MS was combined with 3 optimized CV parameters on the FAIMS Pro Interface to reduced precursor ion interference (Schweppe *et al*, 2019). Data-dependent acquisition (DDA) was performed by selecting the most abundant precursors from each CV’s (-40/-60/-80) MS^1^ scans for hrMS/MS over a 1-1.5s duty cycle (1s/1.5s/1s respectively). The MS^1^ scan parameters include a 400-1,600 m/z mass range at 60,000 resolution (at 200 Th) with 4 × 10^5^ automated gain control (AGC) (100 %), and a maximum injection time (max IT) of 50 ms. Precursors (z=2-5) were isolated with 0.7 Th (quadrupole), fragmented with high energy C-trap dissociation (HCD) at 36 normalized collision energy (NCE), and subjected to hrMS/MS on the Orbitrap at 50,000 resolution (at 200 Th), 120 ms max IT, fixed first mass 110 Th, and 1.0 × 10^5^ AGC (200%). Precursors were placed on 90 s dynamic exclusion (+/-10 ppm) to prevent redundant sampling.

In the iNeuron TMTpro experiments, the same FAIMS and MS^1^ parameters were implemented (with 1.25 s duty cycle/CV for DDA) with the Multi-Notch SPS-MS3 acquisition method (McAlister *et al*, 2014), to further reduce ion interference in TMT reporter quantification (Paulo *et al*., 2016). Most abundant precursors (with 120 s dynamic exclusion +/-10 ppm) were selected from MS^1^ scans, isolated using the quadrupole (0.6 Th isolation), fragmented with collision-induced dissociation (CID) at 35 NCE, and subjected to MS/MS in the ion trap (turbo scan speed, 35 ms max IT, 1.0 × 10^4^ AGC). Using Real Time Search analysis software (Erickson *et al*, 2019; Schweppe *et al*, 2020), a synchronous-precursor-selection (SPS) API-MS^3^ scan was collected on the top 10 most intense b-or y-ions from the matched peptide identification (determined by an online search of its respective MS/MS scan). MS^3^ scans were performed on the Orbitrap (AGC 2.0 × 10^5^; NCE 55; max IT 250 ms, 50,000 resolution at 200 Th). To increase quantitative sampling (MS^3^ scans) of proteins during each mass spectrometry injection, a 2 peptide per protein per sample closeout was set. This ensures no more than two peptide-spectrum matches per protein (that pass quality filters) are subjected to MS^3^ scans, reducing redundant protein MS sampling and potentially increasing proteome depth (Schweppe *et al*., 2020).

### Proteomics Data Analysis

Mass spectrometry raw data were converted to mzXML and monoisotopic peaks were reassigned using Monocle (Rad *et al*, 2021). Mass spectra were database searched using Sequest algorithm (2019.01 rev. 5; (Eng *et al*, 1994)) against the Human Reference Proteome Uniprot database (2019-01 SwissProt entries only; UniProt Constortium, 2015) appended with sequences of common contaminates and reverse sequences of proteins as decoys, for target-decoy competition (Elias & Gygi, 2007). Sequest search parameters include: 50 ppm precursor tolerance, 0.9 Da product ion tolerance, trypsin endopeptidase specificity (C-terminal to [KR], 2 max missed cleavages), static modifications on peptide N-terminus and lysines with TMTpro tags (+304.207 Da) and carbamidomethylation on cysteines (57.021 Da), and variable modification of oxidation on methionines (+15.995 Da). Peptide-spectrum matches were filtered at 2% false discovery rate (FDR) using linear discriminant analysis (Huttlin *et al*, 2010), based on XCorr, DeltaCn, missed cleavages, peptide length, precursor mass accuracy, fraction of matched product ions, charge state, and number of modifications per peptide (additionally restricting PSM Xcorr >1 and peptide length>6). Following a 2% protein FDR filter (Savitski *et al*, 2015), PSMs reporter ion intensities were quantified (most intense centroid within 0.003 Da of theoretical TMT reporter mass), filtered based on a precursor isolation specificity > 0.5, and filtered by a summed signal-to-noise across TMT channels > 100.

Protein quantification was performed via the summation (weighted average) of its constituent PSMs’ reporter intensities and TMT channels were normalized for protein input to total TMT channel intensity across all quantified PSMs (adjusted to median total TMT intensity for the TMT channels) (Plubell *et al*, 2017). Log_2_ normalized summed protein reporter intensities were compared using a Student’s t-test and p-values were corrected for multiple hypotheses using the Benjamini-Hochberg adjustment (Benjamini & Hochberg, 1995). Resultant q-values and mean log_2_ fold changes between conditions were used to generate volcano plots. Hotelling T^2^ analysis was performed using the normalized summed protein reporter ion intensities and the timecourse (v. 1.66.0; (Tai, 2022)) package in R.

The annotation list for the subcellular localization of organellar protein markers was derived from previously published high confidence HeLa dataset ((Itzhak *et al*, 2016); “high” & “very high” confidence) and additional manual entries (Ordureau *et al*., 2021). MitoCharta 3.0 (Rath *et al*, 2021) was used for mitochondrial annotation, whereas annotations for ribosome and autophagy components were used from previous studies (An *et al*, 2020). Transcription factor and neuronal development markers were based on previously published databases and publications (Lambert *et al*, 2018; Ordureau *et al*., 2021). Figures were generated using a combination of Excel, R (v.4.2.0) in RStudio (2022.07.01 Build 554), Perseus (v1.6.5 (Tyanova & Cox, 2018)), GraphPad Prism (v9.1.0), and Adobe Illustrator (26.2.1).

The **Table EV3-8** contains the quantified proteins as well as associated TMT reporter ratio to control channels used for quantitative analysis. The TMT reporter channel assignment used for this study are added to each table.

### Interaction Proteomic Analysis

For interaction analysis of FBXO7 and PINK1, interaction proteomics data from the Bioplex Interactome (Huttlin *et al*., 2021) was extracted for HEK293T and HCT116 samples (**Table EV1 and 2**). The number of replicates in which a peptide-spectral match (PSM) mapped to PINK1’s or FBXO7’s the Bioplex 3.0 interactomes was plotted via a network diagram in R. (packages to add from R include (igraph 1.3.1, tidygraph 1.2.1, and ggraph 2.0.5).

## Microscopy

Macros and pipelines used in this work can be found on GitHub (https://github.com/harperlaboratory/FBXO7.git) or Zenodo (https://zenodo.org/record/7803294).

### Live-cell confocal microscopy for mitophagic flux analysis over differentiation (mt-mKeima^XL^)

For quantitative Keima-flux analysis (Ordureau *et al*., 2020), hESC were seeded on day 4 of differentiation into 6-well 1.5 high performance glass bottom plates (Cellvis, P06-1.5H-N) and further differentiated in the vessel for the indicated times until reaching appropriate confluency and imaged. iNeurons were imaged using a Yokogawa CSU-W1 spinning disk confocal on a Nikon Eclipse Ti-E motorized microscope. The system is equipped with a Tokai Hit stage top incubator and imaging was performed at 37°C, 5% CO_2_ and 95% humidity under a Nikon Plan Apo 60×/1.40 N.A immersion oil objective lens. For ratiometic imaging, mtKeimaXL were excited in sequential manner with a Nikon LUN-F XL solid state laser combiner ([laser line – laser power]: 445 - 80mW, 561-65mW]) using a Semrock Di01-T445/515/561 dichroic mirror. Fluorescence emissions were collected through a Chroma ET605/52m [for 445 nm] and a 568 Chroma ET605/52m [for 561 nm], filters, respectively (Chroma Technologies). Confocal images were acquired with a Hamamatsu ORCA-Fusion BT CMOS camera (6.5 μm^2^ photodiode, 16-bit) camera and NIS-Elements image acquisition software. Consistent laser intensity and exposure time were applied to all the samples, and brightness and contrast were adjusted equally by applying the same minimum and maximum display values in ImageJ/FiJi (Schindelin *et al*., 2012). Image quantification was performed in ImageJ/FiJi using custom-written batch-macros.

In brief, raw confocal images of mitochondrial targeted mt-mKeima^XL^ were divided [ex:561/ex:445] resulting in a ratiometic image of only acidic Keima-puncta. These signals were subjected to background subtraction (rolling kernel size 25, sliding paraboloid) and converted to binary objects. The “Analyze Particles…” command (pixel size exclusion: 0.5-∞, exclude edge objects) was used to measure foci-abundance and other morphological parameters. Results for each image-stack saved as .csv files, together with the original ratiometic .tiff file for QC purposes. Unless stated otherwise all images represent z-projections. Statistical analysis and plotting of microscopy data was performed in Prism (v9.1.0, GraphPad).

### Immunocytochemical analysis

iNeurons or HeLa cells were fixed with warm 4% paraformaldehyde (Electron Microscopy Science, #15710, purified, EM grade) in PBS at 37°C for 30 min and permeabilized with 0.5% Triton X-100 in PBS for 15 minutes at room temperature. After three washes with 0.02% Tween20 in PBS (PBST), cells were blocked for 10 min in 3% BSA-1xPBS at room temperature and washed again three times in PBST. Cells were incubated for 3h in primary antibodies in 3% BSA-1xPBS and washed three times with PBST. Secondary antibodies (Thermo Scientific, 1:400 in 3% BSA-1xPBS) where applied for 1h at room temperature. Alexa Fluor 633 Phalloidin (Thermo Fisher, 1:200, A22284) was added with secondary antibodies to label F-actin. To stain nuclei, Hoechst33342 (1:10000) was added for 5 min to cells in PBST and final three washes performed before mounting in Vectashield (Vector Laboratories, H-1000-10). Primary and secondary antibodies used in this study can be found in the **Materials Table (Table EV9)**.

### Fixed-cell microscopy–general acquisition parameters

Immunofluorescently labelled Hela or iNeurons (antibodies indicated in figures and figure legends and details in **Materials Table, (Table EV9)**) were imaged at room temperature using a Yokogawa CSU-W1 spinning disk confocal on a Nikon Eclipse Ti-E motorized microscope equipped with a Nikon Plan Apochromat 40×/0.40 N.A air-objective lens, Nikon Apochromat 60×/1.42 N.A oil-objective lens and a Plan Apochromat 100×/1.45 N.A oil-objective lens. Signals of 405/488/568/647 fluorophores were excited in sequential manner with a Nikon LUN-F XL solid state laser combiner ([laser line – laser power]: 405 - 80mW, 488 - 80mW, 561 - 65mW, 640nm - 60mW]) using a Semrock Di01-T405/488/568/647 dichroic mirror. Fluorescence emissions were collected with Chroma ET455/50m [405 nm], 488 Chroma ET525/50m [488 nm], 568 Chroma ET605/52m [561 nm], 633 Chroma ET705/72m [640 nm] filters, respectively (Chroma Technologies). Confocal images were acquired with a Hamamatsu ORCA-Fusion BT CMOS camera (6.5 μm^2^ photodiode, 16-bit) camera and NIS-Elements image acquisition software. Consistent laser intensity and exposure time were applied to all the samples, and brightness and contrast were adjusted equally by applying the same minimum and maximum display values in ImageJ/FiJi (Schindelin *et al*., 2012).

### Microscopy-based mitochondrial morphology measurements in iNeurons

For quantitative measurement of mitochondrial morphology in d12 iNeurons under fed or mitophagy conditions, HeLa cells were seeded on 6-well 1.5 high performance glass bottom plates (Cellvis, P06-1.5H-N) and mitophagy induced for the indicated time durations. Alternatively, hESC were seeded on day 4 of differentiation into 6-well 1.5 high performance glass bottom plates (Cellvis, P06-1.5H-N) and differentiated in the vessel into iNeurons and mitophagy induced for the indicated time durations. Cells were fixed and stained as described above. Z-stacks were acquired with a Nikon Plan Apo 100×/1.45 N.A oil-objective lens and with the parameters stated above. Image quantification was performed in ImageJ/FiJi using custom-written batch-macros.

Statistical analysis and plotting of microscopy data was performed in Prism (v9.1.0, GraphPad). Primary and secondary antibodies used in this study can be found in the **Materials Table (Table EV9)**.

### Microscopy-based mtDNA turnover measurements in HeLa and iNeurons

For quantitative measurement of mtDNA turnover after AO-induced mitophagy, HeLa cells were seeded on 6-well 1.5 high performance glass bottom plates (Cellvis, P06-1.5H-N) and mitophagy induced for the indicated time durations. Alternatively, hESC were seeded on day 4 of differentiation into 6-well 1.5 high performance glass bottom plates (Cellvis, P06-1.5H-N) and differentiated in the vessel into iNeurons and mitophagy induced for the indicated time durations. Cells were fixed and stained as described above, but primary antibody incubation with aDNA (1:200) was performed overnight. Z-stacks were acquired with a Nikon Plan Apo 100×/1.45 N.A oil-objective lens and with the parameters stated above. Image quantification was performed in ImageJ/FiJi using custom-written batch-macros. In brief, both aDNA and nuclear signals were converted to binary files and the nuclear signal subtracted from the aDNA signal, resulting in an image stack containing only the mtDNA intensities. The “Analyze Particles…” command (pixel size exclusion: 0.05-3, exclude edge objects) was used to measure morphological features and results for each image-stack saved as .csv files, together with the analyzed binary-mask overlay .tiff file for QC purposes. Number of mtDNA signals were normalized to cell number found in the same image stack. Statistical analysis and plotting of microscopy data was performed in Prism (v9.1.0, GraphPad). Primary and secondary antibodies used in this study can be found in the **Materials Table (Table EV9)**.

### Microscopy-based measurements of p62 recruitment in HeLa

For quantitative measurement of p62 recruitment to mitochondria, HeLa control and knockout cells were seed in 1.5 high performance glass bottom plates and doxycycline added over night to induce Parkin expression. Mitophagy was induced using Oligomycin A / Antimycin A for 16h in full DMEM in presence of doxycycline. Cells were fixed and stained as stated above and imaged using a Nikon Plan Apo 60×/1.42 N.A air-objective lens and with the parameters stated above. 8 μM z-stacks were taken for each selected field of view.

Image quantification was performed in ImageJ/FiJi using custom-written batch-macros. The p62 channel was filtered (Gaussian Blue, sigma =2) and converted into binary files using the Intermodes thresholding method. p62 spots were counted using the “Analyze Particles…” function (pizel size exclusion: 0.1-30, exclude edge objects) and results for each image-stack saved as .csv files, together with the analyzed binary-mask overlay .tiff file for QC purposes. The DAPI channel was used to count nuclei for per cell normalization. Statistical analysis and plotting of microscopy data was performed in Prism (v9.1.0, GraphPad). Primary and secondary antibodies used in this study can be found in the **Materials Table (Table EV9)**.

### Microscopy-based pUb-coverage measurements of mitochondria in iNeurons

For quantitative measurement of pUb coverage over mitochondria AO-induced mitophagy, hESC were seeded on day 4 of differentiation into 6-well 1.5 high performance glass bottom plates (Cellvis, P06-1.5H-N) and differentiated in the vessel into iNeurons and mitophagy induced for the indicated time durations. Cells were fixed and stained as described above, using anti-HSP60 to label mitochondria and anti-pUb (Ser65) to label pUb. 3-5 10 μm thick z-stacks per replicate per sample were acquired using a Nikon Plan Apo 60×/1.42 N.A air-objective lens and with the parameters stated above.

Image quantification was performed in ImageJ/FiJi using custom-written batch-macros. In brief, mitochondrial signal was filtered (Gaussian Blur, sigma=2), converted into binary files and holes in the resulting mask filled. pUb channel was thresholded into a binary file (Triangle method); these masks were measured using the “Analyze Particles…” command (pixel size exclusion: 0.5-**∞**, exclude edge objects) and results for each image-stack saved as .csv files, together with the analyzed binary-mask overlay .tiff file for QC purposes. % of mitochondrial pUb coverage was calculated and normalized to [t]=6h AO. Statistical analysis and plotting of microscopy data was performed in Prism (v9.1.0, GraphPad). Primary and secondary antibodies used in this study can be found in the **Materials Table (Table EV9)**.

### Microscopy-based evaluation of Parkin translocation and mitophagy in FBXO7^-/-^ cell lines

For quantitative measurement of Parkin translocation kinetics in FBXO7^-/-^ cell lines, stable HeLa or HEK293T cell lines expressing GFP-Parkin were created. WT and knockout cell lines were seeded into 12, 24 or 96-well 1.5 high performance glass bottom plates (Cellvis, P12/24/96-1.5H-N) two days prior to experimental manipulation. Mitophagy was induced as described above using Antimycin A / Oligomycin A or CCCP for 1h. Treated and control (fed) cells were fixed as described above, stained for mitochondria (HSP60), Parkin (GFP) and DNA (SpyDNA-555) and 8μm z-stacks were acquired with a Nikon Plan Apo 60×/1.42 N.A oil-objective lens and with the parameters stated above. HEK293T cells were stained for mitochondria (HSP60), Parkin (GFP) and DNA (Hoechst) and 8μm z-stacks were acquired with a Nikon Plan Apochromat 40×/0.40 N.A air-objective lens, using the HCA-module in NIS-Elements.

For quantitate single-cell analysis, a total of ∼240 (HeLa) or 135 (HEK293T) randomly xy-marked stacks were acquired, and MIPs used for subsequent analysis. CellProfiler (Stirling *et al*., 2021) was used for the quantitative analysis of single-cells and mitochondrial objects (segmentation and analysis pipelines can be found on GitHub). Plotting of microscopy data was performed in Prism (v9.1.0, GraphPad) and R using the following libraries: tidyverse, dplyr, tibble, viridis, ggplot2, ggridges, ggsci. Primary and secondary antibodies used in this study can be found in the **Materials Table (Table EV9)**.

### Microscopy-based evaluation pUb recruitment after G-TPP treatment in HeLa FBXO7^-/-^ cell lines

For quantitative measurement of pUb recruitment to mitochondria after G-TPP treatment, HeLa cell lines were seeded into 96-well 1.5 high performance glass bottom plates (Cellvis, 96-1.5H-N) two days prior to experimental manipulation. Cells were treated for the indicated times and fixed and immunofluorescently labelled as described above. Cells were stained for pUb, Parkin (GFP) and DNA (Hoechst33342) and 8μm z-stacks were acquired with a Nikon Plan Apochromat 40×/0.40 N.A air-objective lens, using the HCA-module in NIS-Elements.

9 stacks of randomly selected field of views were taken and MIPs used for analysis using CellProfiler. Plotting of microscopy data was performed in Prism (v9.1.0, GraphPad). Primary and secondary antibodies used in this study can be found in the **Materials Table (Table EV9)**.

### Immunocytochemical sample preparation for 3D-SIM

Sample preparation and SIM acquisition guidelines from (Kraus *et al*, 2017) were used as a basis for the super-resolution analysis of pUb spreading after mitophagy induction. iNeurons or HeLa cells were seeded on 18×18 mm Marienfeld Precision cover glasses thickness No. 1.5H (tol. ± 5 μm). Cover glasses were coated, if necessary, for hESC / iNeuron culture as described above. After experimental manipulation, cells were fixed with warm paraformaldehyde 3% Glutaraldehyde 0.35% in 0.1M Sodium Cacodylate, pH 7.4 (Electron Microscopy Science) at 37°C for 30 min and permeabilized with 0.5% Triton X-100 in PBS for 15 minutes at room temperature. After three washes with 0.02% Tween20 in PBS (PBST), cells were blocked for 10 min in 3% BSA-1xPBS at room temperature and washed again three times in PBST. iNeurons were incubated overnight in primary antibodies in 3% BSA-1xPBS and washed three times with PBST. Secondary antibodies (Thermo Scientific, 1:400 in 3% BSA-1xPBS) where applied for 1 h at room temperature. To stain nuclei, DAPI was added for 5 min to cells in PBST, washed three times for 5 min in 1xPBST before a post-fixation in 4% paraformaldehyde was performed. After 2 washes in PBST, coverslips were washed once in 1xPBS and mounted in Vectashield (Vector Laboratories, H-1000-10) on glass slides. Primary and secondary antibodies used in this study can be found in the **Materials Table (Table EV9)**.

### 3D-SIM microscopy–acquisition parameters

3D-SIM microscopy was performed on a DeltaVision OMX v4 using an Olympus 60x / 1.42 Plan Apo oil objective (Olympus, Japan). The instrument is equipped with 405 nm, 445 nm, 488 nm, 514 nm, 568 nm and 642 nm laser lines (all >= 100 mW) and images were recorded on a front-illuminated sCMOS (PCO Photonics, USA) in 512×512px image size mode, 1x binning, 125 nm z-stepping and with 15 raw images taken per z-plane (5 phase-shifts, 3 angles). Raw image data was computationally reconstructed using CUDA-accelerated 3D-SIM reconstruction code (https://github.com/scopetools/cudasirecon) based on (Gustafsson *et al*, 2008). Optimal optical transfer function (OTF) was determined via an in-house build software, developed by Talley Lambert from the NIC / CBMF (GitHub: https://github.com/tlambert03/otfsearch, all channels were registered to the 528nm output channel, Wiener filter: 0.002, background: 90).

### 3D-SIM microscopy– pUb-mitochondria 3D renderings & analysis

3D renderings of 3D-SIM images were performed in UCSF Chimera X (https://www.cgl.ucsf.edu/chimerax/) using 32-bit .tiff image stacks. Image channels were sequentially imported and visualized as surfaces using the “Volume Viewer” tool. Surface representations were cleaned up using the “High dust” command, based on size filtering thresholding.

3D-analysis of HeLa and iNeuron datasets from 3D-SIM datasets was performed using Imaris (Oxford Instruments, v9.7). After converting all multi-color .tiff to native .ims files, import into Imaris Arena and global background subtraction, mitochondrial and pUb objects were segmented from seeds (XY starting diameter: 0.08 μm == pixel size of images), segmented based on automatic thresholding with local background subtraction and splitting of touching objects (0.4μm). Objects were piped into Imaris Vantage module for further analysis. In Vantage, nearest neighbour distances of pUb to pUb and between pUb and mitochondria, as well as volume of segmented objects were computed. This pipeline was tested on WT control cells and then applied for batch processing on all other genotype to allow for unbiased segmentation and analysis. On average, 10 cells were analysed per genotype per condition (121 total) for HeLa cells, and 12 cells (139 total) for iNeurons.

Unless stated otherwise, all images depicted in figures are maximum-intensity projections.

### Flow cytometry-based measurement of mitophagic flux

HeLa HFT/TO-PRKN cells of the indicated genotype (control, PINK1^-/-,^ FBXO7 ^-/-^) expressing mtKeima (Heo *et al*., 2015) were seeded into 12-well dishes. Upon reaching 60% confluency, 2 μg/ml doxycycline was added to induce Parkin expression for at least eight hours before depolarization of mitochondria. Following Parkin induction, cells were treated with Antimycin A (5 μM) and Oligomycin (10 μM) for the indicated number of hours. Bafilomycin A (25 nM, Sigma-Aldrich, B1793)-treated samples were treated for three hours. Control cells were fed three hours before experimental manipulation. To harvest cells for analysis, each well was washed with 1 mL PBS then treated with 100 μL of 0.25% trypsin for 3 min at room temperature, resuspended in 300 μL of DMEM with 10% FBS. 200 μL of each sample was transferred to a flat-bottom 96-well plate for analysis by flow cytometry.

hESCs were seeded at day 4 into coated 6-well dishes and differentiated. At day 12, mitophagy was induced with Antimycin A (5 μM) and Oligomycin (10 μM) or Bafilomycin A (25 nM, Sigma-Aldrich, B1793) for the indicated number of hours. Control cells were fed 2h before experimental manipulation. Cells were dissociated from the wells using Accutase and resuspended in 600 μL ND2 medium + 10 μM Y27632 and filtered through a cell strainer cap tube (Corning, 352235).

Analysis of a population of at least 10,000 cells was performed on an Attune NxT (Thermo Fisher Scientific) detecting neutral mtKeima signal with excitation at 445 nm and emission 603 nm with a 48 nm bandpass and acidic mtKeima with 561 nm excitation and emission 620 nm and a 15 nm band pass. mtKeima ratio was analysed as previously described (An & Harper, 2018). Briefly, acidic:neutral Keima ratios were measured by gating a population of well-behaved single cells using forward and side scatter, followed by calculation of acidic:neutral mtKeima ratio on a per-cell basis in FlowJo Software (FlowJo, LLC).

## Quantification and Statistical Analysis

Unless stated otherwise all quantitative experiments were performed in triplicate and average with S.E.M. or S.D. as indicated in legends reported. All numerical data used in plots can be found in **Table EV10**.

