## Appendix Figure S1-S5 for "PARK15/FBXO7 is dispensable for PINK1/Parkin mitophagy in iNeurons and HeLa cell systems"

**Table of Contents:**

Appendix Figure S1 p.2
Appendix Figure S2 p.4
Appendix Figure S3 p.5
Appendix Figure S4 p.7
Appendix Figure S5 p.8

### Appendix Figure S1

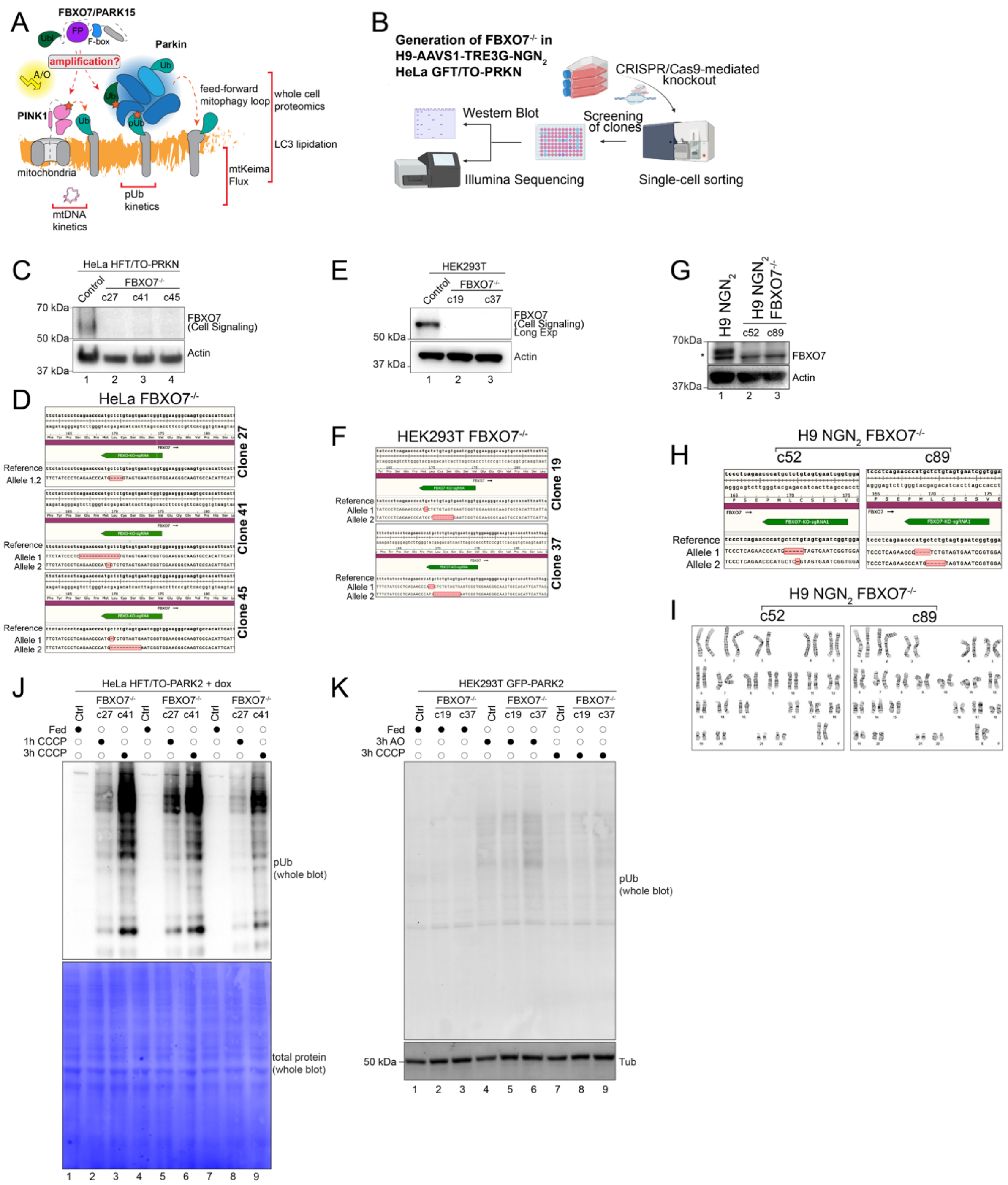


**Appendix Figure S1: Tool kit for analysis of HeLa cells and iNeurons lacking FBXO7.** (**A**) Working model summarizing the suggested modes of action of FBXO7 in PINK1/Parkin mitophagy. (**B**) gRNA sequence within FBXO7 gene along with the sequences of alleles 1 and 2. MiSeq analysis of the three clones used for this study are shown. (**B**) Schematic for generation of FBXO7$^{-/-}$ in HeLa and hESC cell lines. (**C**) Western Blot analysis on HeLa control and FBXO7$^{-/-}$ whole cell lysate. (**D**) Targeting of gRNA sequence within FBXO7 gene along with the sequences of alleles A and B. MiSeq analysis of the three HeLa clones (c27, c41, c45) used for this study are shown. (**E**) Western Blot analysis of HEK293T control and FBXO7$^{-/-}$ whole cell lysate. (**F**) MiSeq analysis of the two HEK293T clones (c19 and c37). (**G**) Western Blot analysis on hESC control and FBXO7$^{-/-}$ whole cell lysate. Asterisk indicates non-specific band. (**H**) Targeting of gRNA sequence within FBXO7 gene along with the sequences of alleles A and B. MiSeq analysis of the two clones (c52, c89) used for this study are shown. (**I**) Karyotype analysis of FBXO7$^{-/-}$ c52 and c89 hESCs. (**J**) Immunoblot of HeLa Control and FBXO7$^{-/-}$ cells after mitophagy induction using CCCP for indicated times. (**K**) Immunoblot of HEK293T Control and FBXO7$^{-/-}$ (expressing GFP-Parkin) after mitophagy induction after indicated times.

### Appendix Figure S2

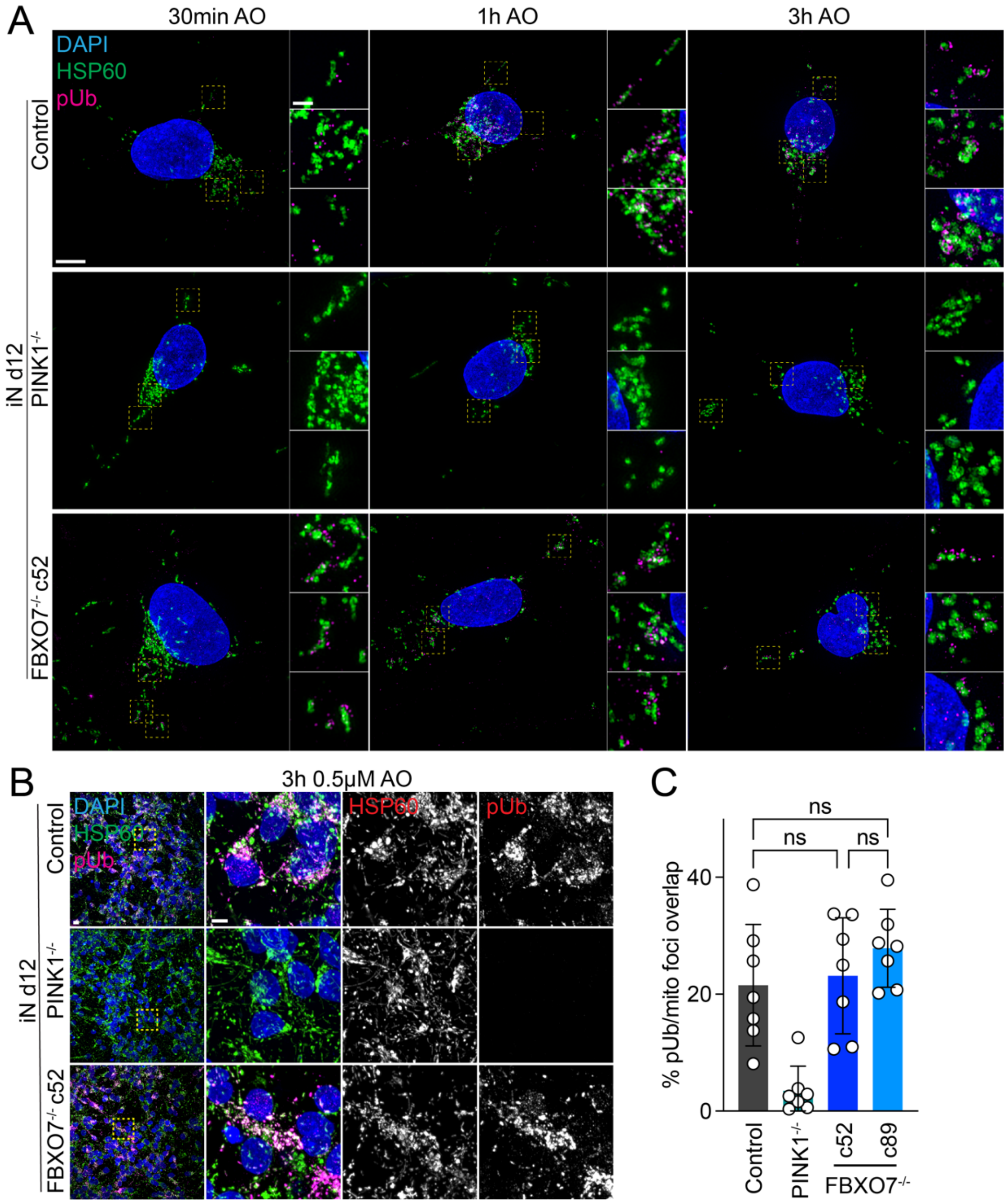


**Appendix Figure S2. Super-resolution pUb detection in iNeurons in response to mitochondrial depolarization.** (**A**) 3D-SIM images of iNeurons day 12 control, PINK1$^{-/-}$ and FBXO7$^{-/-}$ cell lines after AO-induced mitophagy. Cells were stained for nuclear DNA (DAPI), mitochondria (HSP60) and pUb. Related to **Figure 2D**. Scale bar = 5 μm or 1μm. (**B**) Confocal images of iNeurons day 12 control, PINK1$^{-/-}$ and FBXO7$^{-/-}$ cell lines after AO-induced mitophagy with lower AO concentrations. Cells were stained for nuclear DNA (DAPI), mitochondria (HSP60) and pUb. (**C**) Evaluation of images depicted in **B.** Error bars depict S.D. from 12 image stacks measured per condition. One-way ANOVA with multiple comparisons. Scale bar = 10 μm and 5 μm.

### Appendix Figure S3

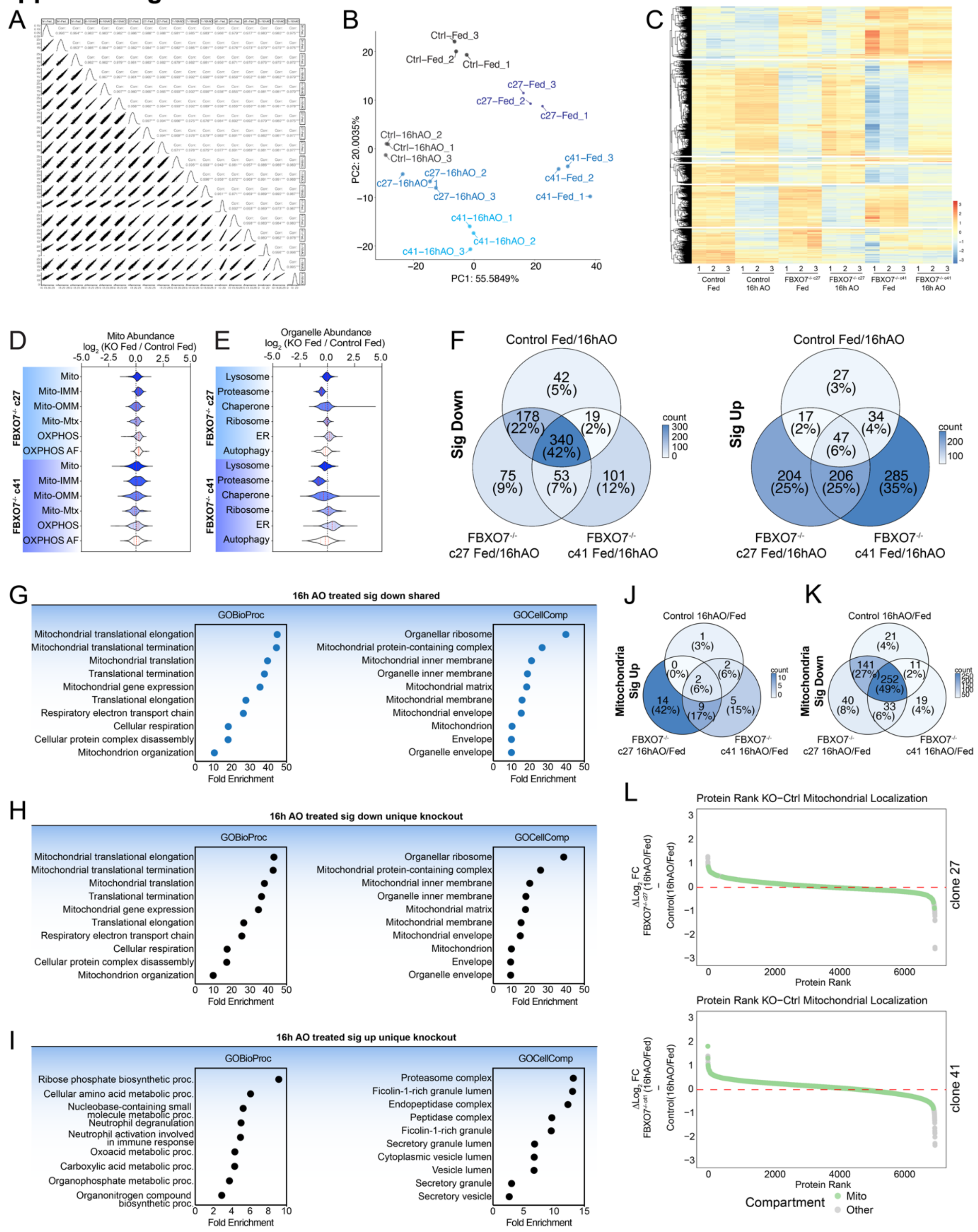


**Appendix Figure S3. Proteomic analysis of HeLa cells lacking FBXO7 during mitophagy.** (**A,B**) Correlation plots (panel A) and PCA analysis (panel B) for ~8000 proteins quantified by TMT proteomics in individual replicates for the experiment outlined in **Figure 4A**. (**C**) Hierarchical clustering of 8000 proteins quantified in the experiment outlined in **Figure 4A**. (**D,E**) Violin plots [$Log_2$ (control / FBXO7$^{-/-}$)] for FBXO7$^{-/-}$ c27 and c41 cells depicting alterations in the abundance of mitochondrial proteins (left panel) or specific organelles or protein complexes (right panel). (**F**) Venn diagrams depicting the overlap in the number of proteins whose levels are reduced (left panel) or increased (right panel) for the data shown in **Figure 4B** are shown. Proteins whose abundance is significantly reduced upon AO treatment are enriched for the mitochondrial organelle (right panel), as expected for cells undergoing mitophagy. (**G**) GO-term enrichment analysis of genes significantly down in both Control and FBXO7$^{-/-}$ after 16h AO mitophagy. (**H,I**) GO-term enrichment analysis of unique genes significantly down (panel H) or up (panel I) in FBXO7$^{-/-}$ after 16h AO mitophagy. (**J,K**) Venn diagrams depicting the overlap in the number of mitochondrial proteins whose levels are significantly increased (**J**) or decreased (**K**) are shown. The plurality of proteins which are significantly decreased, are shared in all cell lines. (**L**) Ranked protein abundance of $\Delta$ log2 (KO[16h AO/Fed] – Control [16h AO/Fed]) for FBXO7$^{-/-}$ clone 27 (top) and clone 41 (bottom). Mitochondrial annotated proteins are depicted green, all other detected proteins in grey.

### Appendix Figure S4

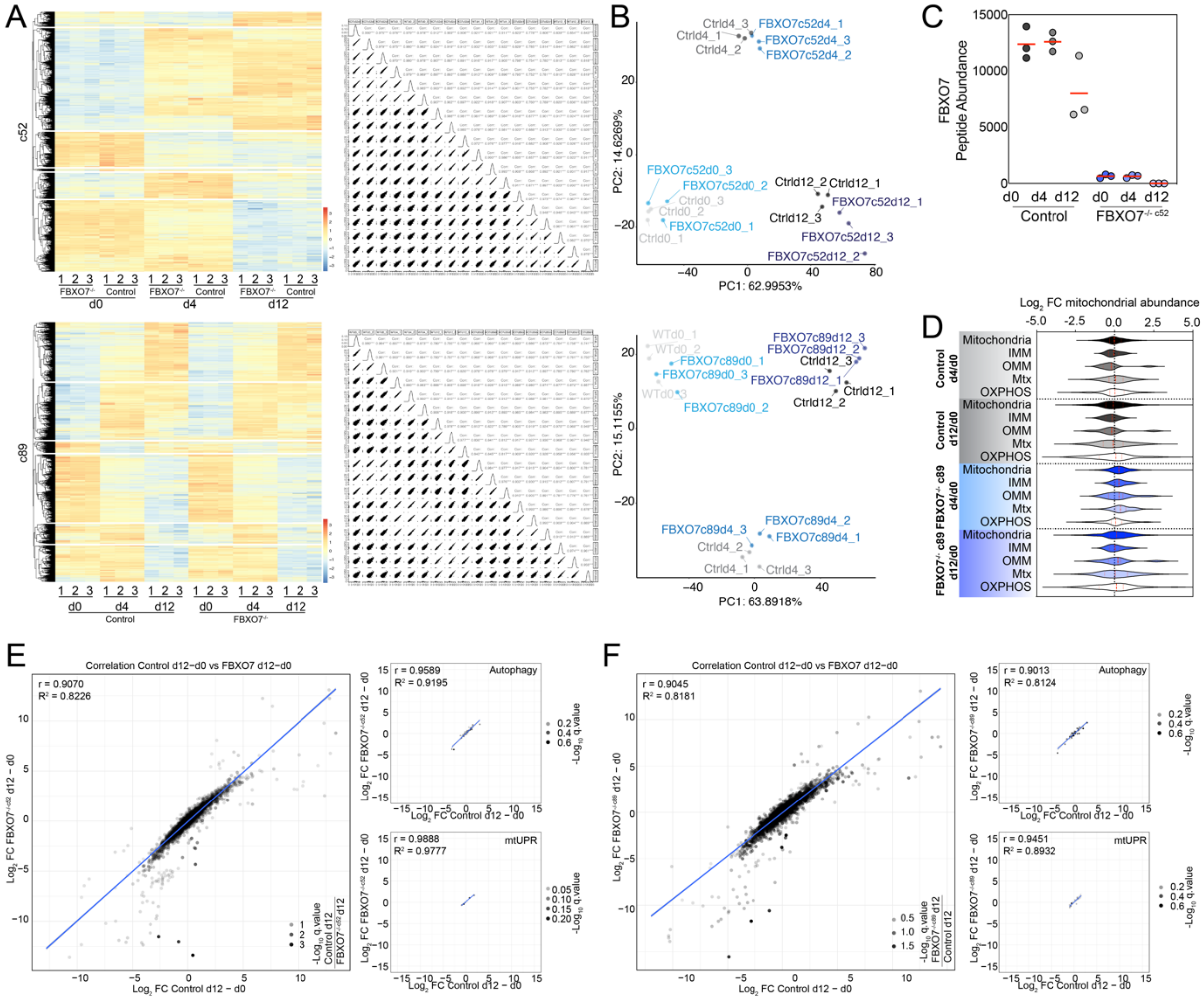


**Appendix Figure S4. Proteomic analysis of human ES cells lacking FBXO7 during neurogenesis.** (**A**) Hierarchical clustering and correlation plots of control and FBXO7$^{-/-}$ c52 (top) or c89 (bottom) at indicated days of differentiation in the experiment outlined in **Figure 5A.** (**B**) PCA analysis of whole cell proteomic samples from control and FBXO7$^{-/-}$ cells. Triplicates for each timepoint and genotype were used. (**C**) FBXO7 peptide abundance in control and FBXO7$^{-/-}$ cells at indicated days of differentiation. (**D**) Violin plots [$Log_2$ (control / FBXO7$^{-/-}$)] for FBXO7$^{-/-}$ c89 cells depicting alterations in the abundance of mitochondrial proteins. (**E,F**) Correlation plot of d12 / d0 of Control vs FBXO7$^{-/-}$ cells (**E**: clone 52; **F**: clone89). Autophagy and mtUPR proteis are plotted on the right. Pearson's r correlation and $R^2$ of KO vs Ctrl are indicated on the top left corners.

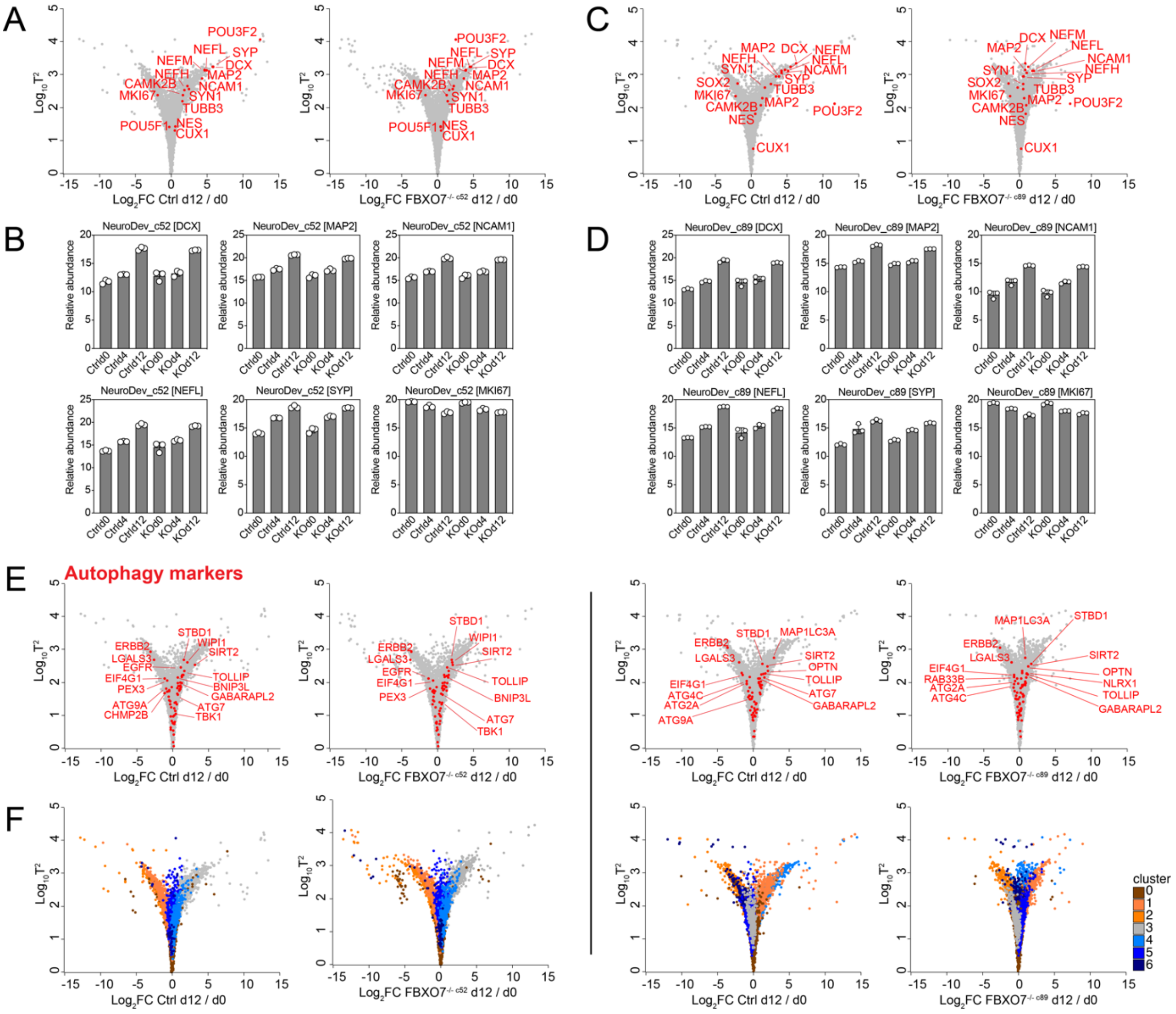


**Appendix Figure S5. Comparison of human ES cell neurogenesis with or without FBXO7.** (**A**) Hotelling plots ($Log_{10}$ $T^2$ statistic versus $Log_2$ FC day 12/day 0) of control (left panel) or FBXO7$^{-/-}$ c52 hES cells undergoing differentiation. Selected neurogenesis factors are indicated. (**B**) The relative abundance of selected proteins is shown in the lower histograms at day 0, 4 and 12 of differentiation. (**C,D**) Hotelling plots similar to **A,B** for control and FBXO7$^{-/-}$ c89. (**E**) Patterns of changes in the abundance of autophagy proteins in control (left panel) and FBXO7$^{-/-}$ (right panel) day 12 iNeurons relative to day 0 cells displayed on a Hotelling plot. (**F**) Cluster analysis of control and FBXO7$^{-/-}$ cells comparing day 12 of differentiation to day 0. Individual clusters are displayed on Hotelling plots for each genotype.
